# Genomic locus of lncRNA-*Gm26793* forms an inter-chromosomal molecular lock with *Cubn* to ensure proper stem cell differentiation and mouse embryogenesis

**DOI:** 10.1101/2023.09.13.557495

**Authors:** Zhiwen Liu, Xianfa Yang, Jiehui Chen, Yongjian Ma, Xing Wan, Yonggao Fu, Yingying Chen, Mingzhu Wen, Yun Qian, Yong Zhang, Dahai Zhu, Jinsong Li, Naihe Jing

**Author notes:** These authors contributed equally (Zhiwen Liu, Xianfa Yang, Jiehui Chen, Yongjian Ma, Xing Wan, Yonggao Fu). Correspondence (X.Y.), (D.Z.), (J.L.), (N.J.).

## Abstract

Inter-chromosomal interactions play a crucial role in 3D genome organization, yet the organizational principles and functional significances remain elusive. In general, long non-coding RNA (lncRNA) loci and transcripts are frequently associated with transcriptional programs modulated by long-range chromatin interactions. Here, we identified a novel lncRNA named *Gm26793*, which is abundantly distributed in the primitive streak and mesodermal cells of E7.5 mouse gastrula. Through genetic ablation of *Gm26793*, we observed a preferential responsiveness to primitive endoderm lineage during stem cell differentiation, as well as enhanced occurrence of transient and degenerative state cells in early mouse embryos when the cell fate segregates between epiblast and primitive endoderm. Mechanistically, we revealed the genomic locus of *Gm26793*, rather than the lncRNA transcript or adjacent gene governs the cell fate preference towards primitive endoderm. Concretely, *Gm26793* locus (Chr 7) forms an inter-chromosomal molecular lock with *Cubn* (Chr 2), restraining the expression of *Cubn* and maintaining a natural epigenetic landscape, thus ensuring the proper lineage specification *in vitro* and *in vivo*. In order to reinforce this lock, CTCF and cohesin complex serves as a ring to fasten the inter-chromosomal contact. Overall, our study provides a clear paradigm that inter-chromosomal interaction collaborates with architectural factors to stabilize nuclear conformation and guarantee faithful gene expression during stem cell differentiation and mammalian embryogenesis.

## Introduction

The mammalian genome encodes tens of thousands of non-coding dark matters, such as long non-coding RNA (lncRNA) genes, which have been found to execute crucial biological functions ^1–3^. However, the precise determination of specific functional lncRNA usually tends to be blind and labor-intensive until the coming era of high-throughput sequencing and efficient genomic editing. Pre-screening through exploration of lncRNA abundance in specific biological tissues, especially across spatial-temporal embryo developmental transcriptomic atlas or disease-related transcriptome reference, could largely facilitate the identification of vital lncRNAs with biological significances ^4–7^. Pioneering mechanistic studies have reported that the lncRNA genes can modulate chromatin structures and regulate the expression of local or distal genes, frequently through the act of transcription and genomic loci ^8–12^. Nevertheless, massive gaps still exist in understanding of the regulatory purposes of ubiquitous lncRNAs and how they differ from the established regulatory network mediated by coding genes. Furthermore, the extent to which genetic removal of diverse lncRNAs can result in physiologically relevant phenotypes remains unclear.

In mammalian cells, the linear sequence of genome is hierarchically organized into distinct chromosome territories (CTs), A/B compartments, topologically associating domains (TADs) and chromatin loops ^13–16^. These structural units ensure the overall genome stability as well as maintain relative plasticity of chromatin interaction against specific physiological stimuli ^17–21^. Among these structural units, the chromatin loops formed via chromatin folding seem to be of the highest flexibility and usually exhibit dramatic dynamics of loop switch upon the stimuli of certain differentiation signals or chemical treatment ^22–25^. As revealed by the typical chromatin interaction capture technology, most of the chromatin loops seem to exist between *cis*-acting anchor sites within merely one chromosome ^26–29^. However, for most biological processes, the genome set usually acts as an entirety and exhibits a coordinated change of conformation upon extracellular stimulation. Thus, how the cells can harmonize the entire set of chromosomes within the nuclei remains largely unknown ^30–34^. The occurrence of direct inter-chromosomal interactions or *trans*-acting contacts provides one potential strategy for the coordination of chromatin conformation in response to certain stimuli ^35–40^. For example, the complex choreography of olfactory receptor genes, which are located across several different chromosomes, involves frequent inter-chromosomal interactions in the form of the “olfactosome” to determine specific olfactory receptor genes’ expression in sensory neurons ^41, 42^. However, whether the inter-chromosomal interactions also exist and execute critical biological functions during early development remains largely unexplored.

The CCCTC-binding factor, CTCF, is a major organizer in orchestrating the chromatin interactions within individual chromosome and between different chromosomes ^43–45^. In the prevailing model of loop-extrusion, cohesin complex can form a “ring” to capture a chromatin loop and slide through the chromatin until they encounter a pair of convergent CTCF binding sites ^46–48^. The boundary areas of chromatin loops within individual chromosomes are generally found to be crucial regulatory elements (such as enhancers, promoters, silencers, and insulators), which are tightly related to gene expression regulation ^49–54^. Therefore, the binding of CTCF engages and determines the coordination of distinct genomic loci, thus ensures the normal gene expression landscape and provides a proper cellular homeostasis ^55–58^. Emerging studies implicate that CTCF-cohesin complex participates in the formation of inter-chromosomal contacts ^59–61^, whereas the specific functional properties of CTCF-cohesin complex in regulating inter-chromosomal interaction await interpretation.

In this study, through systematic analyses of the established mouse spatial transcriptome atlas, we identified a lncRNA gene-*Gm26793*, specifically expressed in the primitive streak and mesoderm tissues of the E7.5 embryo, and found that genetic elimination of *Gm26793* leads to the aberrant upregulation of primitive endoderm genes *in vitro*, and causes developmental arrest during the cell fate segregation between epiblast and primitive endoderm *in vivo*. Molecularly, *Gm26793* (Chr 7) could form inter-chromosomal interaction with *Cubn* (Chr 2) through genomic locus, but independent of the transcript and adjacent gene. This specific inter-chromosomal interaction functions as a molecular lock that limits the expression of *Cubn*, sustaining the appropriate epigenetic modification and differentiation capacity of mESCs. Additionally, akin to the secure fastening of a lock ring, the binding of the CTCF-cohesin complex can fix the linkage of chromatin interactions.

## Results

### The identification of functional lncRNA-*Gm26793* based on the spatial transcriptome atlas of the mouse gastrula

In this study, through in-depth analyses of our recently established spatial transcriptomic atlas of mouse gastrula ^62, 63^, we obtained numerous region-specifically expressed lncRNAs in embryo samples ranging from early-streak stage (E6.5), mid-streak stage (E7.0), to late-streak stage (E7.5) (Fig. 1a and Extended Data Fig. 1a-c, Supplementary Table 1). Generally, we found that lncRNAs could form distinct expression patterns in the gastrula (Fig. 1a and Extended Data Fig. 1b-f), which in accordance with the germ layer-related spatial location during mouse gastrulation ^64, 65^. To specify, based on the differentially expressed lncRNAs (DELs) during gastrulation, three gene groups could be identified with Endoderm (End)-specific (G1, G2) or Epiblast (Epi)-specific (G3) distribution (Extended Data Fig. 1b) for E6.5 embryos. As to the E7.0 embryo, along with the emergence of mesoderm tissues, two new gene groups of lncRNAs (G1, G2) can be identified with mesoderm-specific abundance (Extended Data Fig. 1c). When developing to the E7.5 stage, six gene groups (G1, G2, G3, G4, G5, G6) could be identified (Fig. 1a). In contrast to embryos of the E6.5 and E7.0 stages, primitive streak cells (PS) in the E7.5 stage exhibit specific enrichment of certain lncRNAs, which are clustered into G6 group (Fig. 1a). Further investigation of G6 group related lncRNAs revealed that these lncRNAs are highly expressed in PS, but gradually down-regulated in mesoderm tissues (Mes) and largely absent in endoderm tissues (End) and ectoderm tissues (Ect) (Fig. 1a). Given that mesoderm and definitive endoderm cells are mostly derived from primitive streak through epithelial-mesenchymal transition during gastrulation ^66^, genes harbored in the G6 group may implicate potential biological importance during mesoderm and endoderm development.

**Fig 1.**
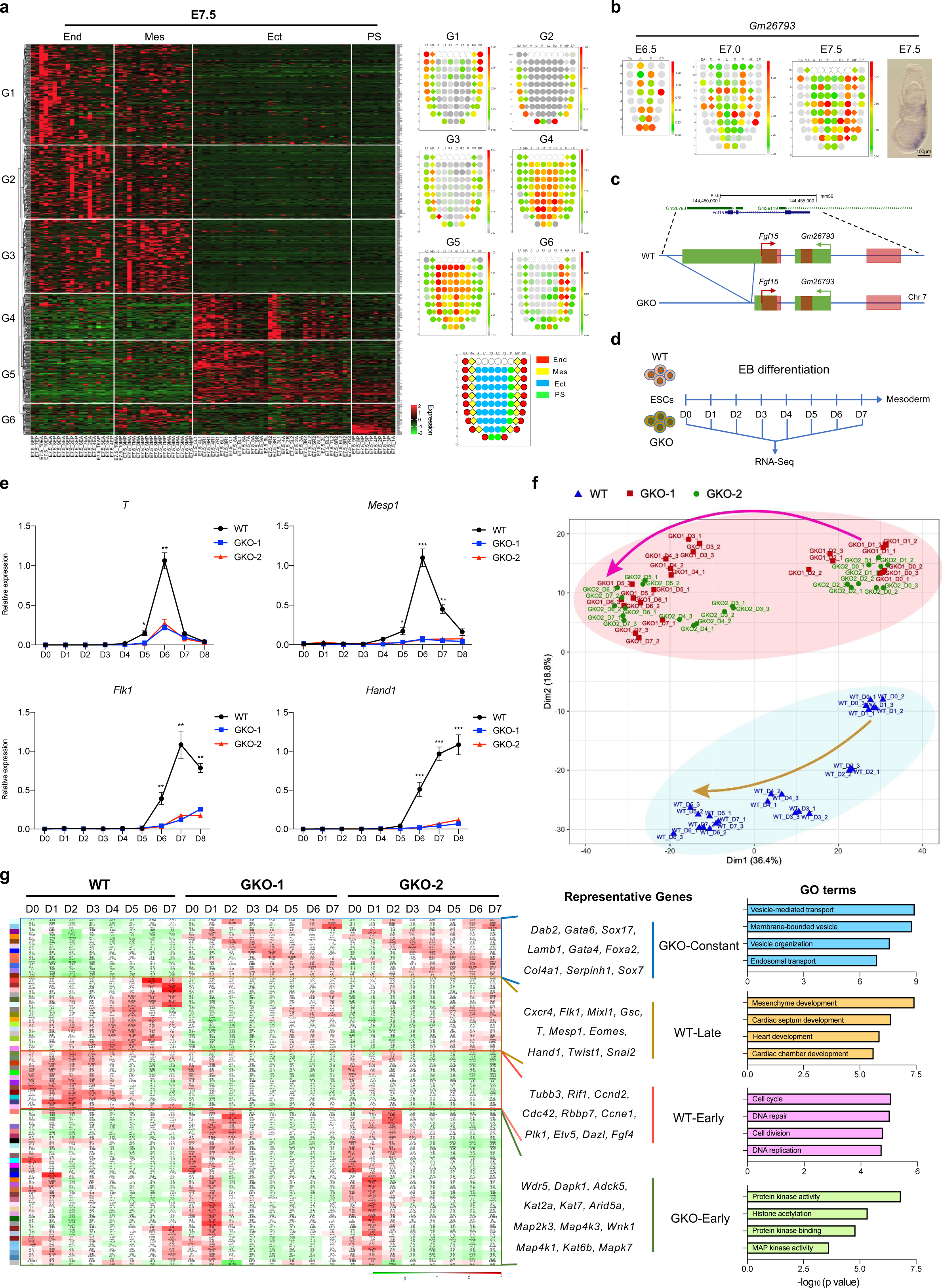
The identification of lncRNA-*Gm26793* distribution in the gastrula and its functional significance during mESCs differentiation. (a) Differentially expressed lncRNAs across Ect, Mes, End and PS regions in the E7.5 embryo. The average expression pattern of each gene group was presented in the right panel as corn plots. End: endoderm; Mes: mesoderm; Ect: ectoderm; PS: primitive streak. (b) The spatial distribution of *Gm26793* transcript in the mouse gastrula. Experimental validation of *Gm26793* expression through whole-mount in situ hybridization was also shown. Scale bar, 100 μm. (c) Schematic diagram showing the strategy for the genomic knockout of *Gm26793*. (d) The workflow depicting the stem cell differentiation program and sampling timepoints for both wild-type and *Gm26793* knockout cells. (e) Time-course expression profiles of mesoderm-related genes during WT and GKO EB differentiation measured by qPCR (n=3). Data were shown as means ± SEM. Student’s *t*-test, \**p*<0.05, ** *p*<0.01, *** *p*<0.001; n represents biologically independent replicates. (f) PCA analyses of EB differentiated samples showing distinct clustering patterns and differentiation trajectories of WT and GKO mESCs. The major differentiation trajectories of both WT and GKO mESCs were highlighted in arrow lines. (g) Heatmap showing stage-specific co-expression gene modules and their correlation to differentiation days. Each row corresponds to a co-expression module. The correlation coefficients and *p*-values are shown in each square. Representative genes and top GO terms of four combined module categories were listed in the right panel.

Amongst the spatial-specific G6 lncRNAs, we found that one lncRNA named as *Gm26793*, which is located adjacently to the protein-coding gene-*Fgf15*, exhibits high enrichment in the PS region of E7.5 embryos (Fig. 1b,c). To investigate the biological significances of *Gm26793*, we took advantage of the CRISPR-Cas9 system to genetically delete the second exon of *Gm26793* (∼1.5 kb) in mouse embryonic stem cells (mESCs), and retain the integrity of *Fgf15* transcripts (Fig. 1c and Extended Data Fig. 2a-c). Two biological replicates with genetic *Gm26793* knockout were prepared, and the resulting knockout embryonic stem cells were named as GKO cells. Pre-examination of the pluripotent marker expression as well as cellular morphology in GKO cells revealed that GKO cells still maintain comparable expression levels of key pluripotent markers in mESCs, such as *Oct4*, *Nanog,* and *Klf4* (Extended Data Fig. 2d,e). In addition, we found that the morphology of GKO clones are relative loosely compacted than wild-type mESCs (Extended Data Fig. 2e,f), suggesting the GKO clone may possess differential developmental potencies in comparison with tightly organized wild-type cells.

### The developmental potency towards mesoderm lineage was largely compromised in GKO cells *in vitro*

To determine the developmental potencies of GKO cells, we firstly subjected both WT and GKO cells to spontaneous differentiation in 10% fetal bovine serum (FBS) medium (Fig. 1d and Extended Data Fig. 2g) ^67^. In accordance with *in vivo* transcriptomic data inferred from the gastrulation atlas, the expression level of *Gm26793* was gradually elevated and peaked at D6 of differentiation (Extended Data Fig. 2h), when the cells reached a mesodermal state. Concomitantly, we found the mesodermal cell markers, such as *T* and *Mesp1*, were largely abolished in GKO cells (Fig. 1e). To systematically assess the effects of *Gm26793* knocking-out, we collected the time-series bulk-cell transcriptomic data of both WT and GKO cells during this process (Fig. 1d). By performing principal component analysis (PCA), we found that the transcriptome differences between WT and GKO cells already exist at the embryonic stem cell stage (D0), and gradually widen along with differentiation from D0 to D7 (Fig. 1f). Next, we applied weighted gene co-expression network analysis (WGCNA) ^68^ to evaluate the time-series transcriptomic distinctions between WT and GKO cells during spontaneous embryoid body (EB) formation. As shown in Fig. 1g, four temporal gene module categories with distinct stage- or sample-specific patterns can be identified, which can be named as WT-Early, WT-Late, GKO-Early, and GKO-constant, respectively (Supplementary Table 2). Gene ontology analyses further identified genes in the WT-late category, which were less expressed in GKO samples, were highly related to mesoderm development (Fig. 1g). qPCR analyses and immunostaining reconfirmed the absence of both transcript and protein levels of mesodermal genes in GKO cells during EB differentiation (Fig. 1e and Extended Data Fig. 2i,j). In contrast, endoderm-related genes, such as *Gata6*, *Sox17*, and *Sox7*, and relevant biological processes, were exclusively enriched in the GKO-constant category (Fig. 1g and Extended Data Fig. 3a). Overall, these results indicate that *Gm26793* knockout severely impedes the spontaneous differentiation of mESCs by down-regulating mesodermal genes and up-regulating endodermal genes.

### The developmental capacity towards primitive endoderm fate was abnormally enhanced in GKO cells *in vitro*

To characterize the differentiation phenotype of GKO cells, we analyzed the enrichment of endoderm-related markers in the differentiated EBs. As expected, we did not detect any expression of endoderm markers, such as GATA6 and SOX17, in the WT EBs (Fig. 2a,b and Extended Data Fig. 3a-c). Interestingly, in contrast to the uniform mesodermal cells distribution in WT EBs, EBs acquired in GKO groups exhibit a two-layer concentric circular structure that cells residing in the inner layer are densely structured, while cells staying in the outer layer are loosely organized and show a strong enrichment with endoderm protein signatures (Fig. 2a,b and Extended Data Fig. 3b,c). To further investigate the molecular features of the GKO EBs, we utilized Geo-seq ^69^ to specifically profile the transcriptome of inner and outer layers of GKO EBs at D8 (Fig. 2c), and found that inner and outer cells of GKO group exhibit distinct transcriptomic patterns unlike the indistinguishable cell composition in WT samples (Fig. 2d and Extended Data Fig. 3d). As revealed by the differentially expressed gene analyses (Supplementary Table 3), we found the absence of mesodermal gene expression in both inner and outer layers of GKO EBs, which are highly expressed in WT EBs. Furthermore, the inner cell layer of GKO EBs expresses a higher level of pluripotency-related genes, such as *Prmt5*, *Sox2*, and *Pou5f1* (Fig. 2d). In contrast, cells residing in the outer layer of GKO EBs showed enrichment of endoderm-related genes, such as *Gata6*, *Foxa2*, and *Sox17* (Fig. 2d). These results suggested that GKO cells fail to initiate mesoderm differentiation, but tend to maintain a stem cell state in the inner layer of EBs, and are more likely to adopt an endoderm fate for cells in the outer layer, which are directly exposed to the signal stimulation,.

**Fig 2.**
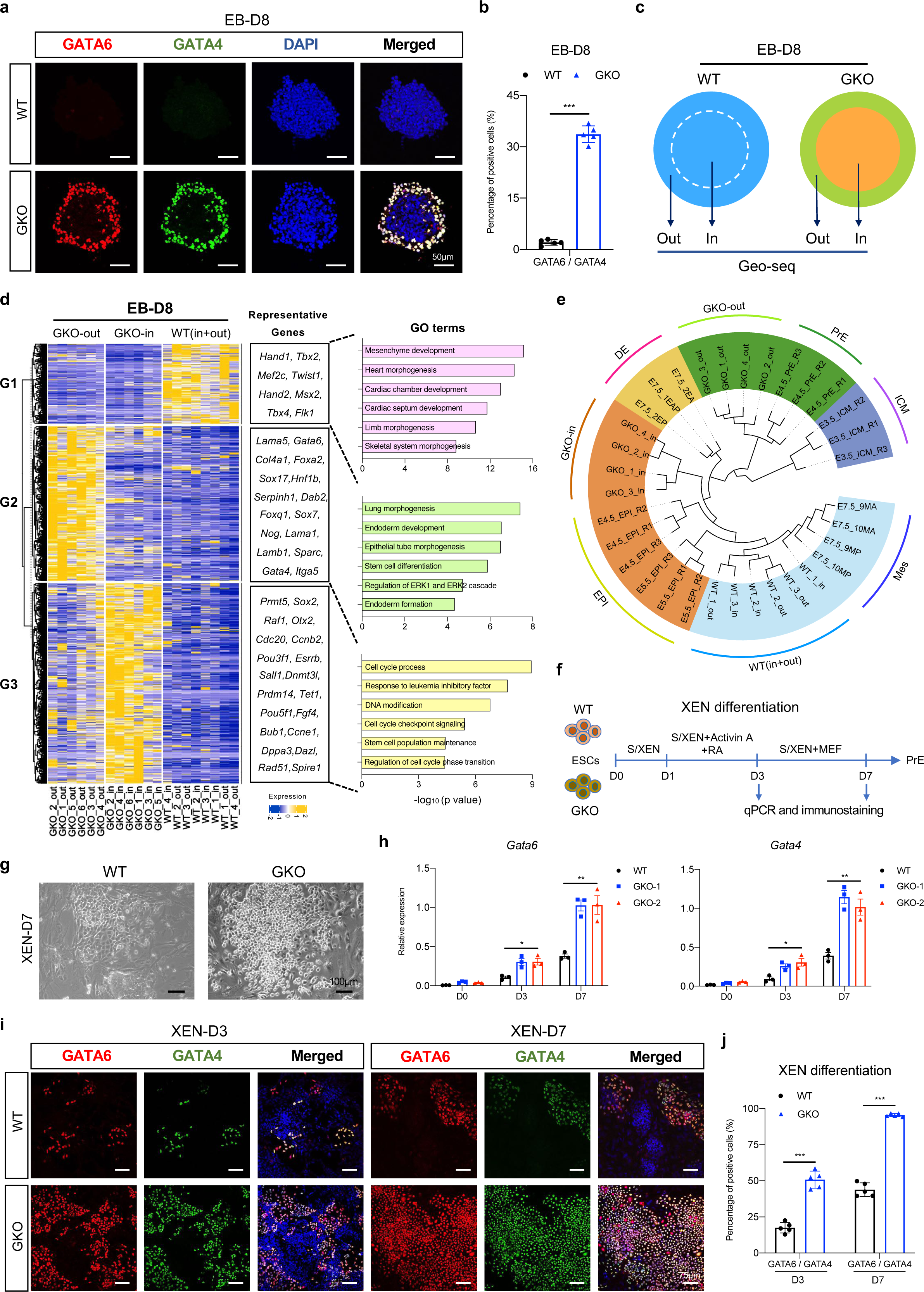
*Gm26793* knockout in mESCs boosts the responsiveness to PrE differentiation signals. (a-b) Immunoflorescence analyses showing the specific distribution of GATA6/GATA4 positive cells in the outer of GKO EBs collected at day 8. Scale bar, 50 μm. Quantification of fluorescent-positive cells was shown in (b). (c) Illustration showing the sample collection of outer and inner layers in WT and GKO EBs by Geo-seq for transcriptomic analysis. (d) Differential expressed gene analyses identify the specific gene expression pattern for each dissected WT and GKO samples. Representative genes and GO terms of each DEG group were listed in the right panel. (e) The phylogenetic tree displaying the clustering of EB differentiation samples with mouse pre-implantation cell lineages as well as definitive endoderm and mesoderm lineages. The mouse pre-implantation, definitive endoderm and mesoderm transcriptomic data are collected from published datasets ^63, 70^. (f) The schematic diagram describing the differentiation protocol for XEN-directed differentiation from embryonic stem cells. (g) The morphologies of differentiated cells in bright field for both WT and GKO XEN cells at day 7. Scale bar, 100 μm. (h) Temporal expression dynamics of *Gata6* and *Gata4* during WT and GKO XEN differentiation. (i-j) Representative images showing the presence of GATA6/GATA4 positive cells in WT and GKO groups at day 3 and 7 during XEN differentiation. Scale bar, 75 μm. Quantification of the data was shown in (j). All data were shown as means ± SEM. Student’s *t*-test, \**p*<0.05, ** *p*<0.01, *** *p*<0.001.

To address the lineage features of the GKO cells, we incorporated *in vivo* published embryogenesis dataset and build a hierarchy of clusters with differentiated EB samples (Fig. 2e). According to the hierarchical clustering results, we found that the inner cells of GKO EBs were clustered with epiblast samples around peri-implantation stages (E4.5_EPI and E5.5_EPI), when epiblast cells still maintain a naïve or formative pluripotent state ^70–72^ (Fig. 2e). Whereas, the outer layer of GKO EBs were closely linked with primitive endoderm samples (E4.5_PrE), but not definitive endoderm samples (E7.5_EA/P) (Fig. 2e). Consistently, we observed that, in contrast to the increased expression of pan-endoderm markers (Fig. 1g and Extended Data Fig. 3a), the expression of mesendodermal makers, *Eomes* and *Gsc* ^73–76^, were severely affected (Extended Data Fig. 3e). Therefore, the increment of pan-endodermal markers’ expression, such as *Gata6* and *Sox17,* should be attributed to elevated primitive endoderm differentiation.

In view of the enhanced responsiveness to primitive endoderm commitment upon *Gm26793* knockout, we explored whether this preference is retained in directed extraembryonic endoderm (XEN) differentiation system (Fig. 2f) ^77, 78^. Along with 7 days of differentiation, we found that nearly all GKO cells turned into highly refractile phase-bright XEN, a typical morphological characteristic of mature XEN cells (Fig. 2g). In contrast, only a subset of WT cells could achieve the epithelial-like XEN state, and the majority of WT cells remain in undifferentiated compact status (Fig. 2g). qPCR ananlysis showed the expression levels of primitive endoderm markers, such as *Gata6*, *Gata4*, *Sox17*, *Foxa2*, *Sox7*, and *Dab2*, were gradually up-regulated in both WT and GKO cells (Fig. 2h and Extended Data Fig. 3f). But, the increment of primitive endoderm markers in GKO cells was much faster than in WT cells. Similarly, immunostaining results also revealed the protein signatures of primitive endoderm were markedly elevated in GKO cells (Fig. 2i,j and Extended Data Fig. 3g,h), which indicates that *Gm26793* knockout indeed boosts the differentiation of mESCs towards primitive endoderm lineage.

### *Gm26793* null embryos exhibit developmental failure during early lineage segregation between epiblast and primitive endoderm *in vivo*

In order to determine the roles of *Gm26793* during mouse embryogenesis, we generated *Gm26793* knockout mice by removing the same genomic region as GKO cells (Extended Data Fig. 4a). Generally, mice without *Gm26793* were viable. However, after summarizing the genotype of mice acquired from heterozygous parents, we found a non-negligible consistent loss of homozygous offspring (Extended Data Fig. 4b), which implied a portion of homozygous GKO mice could be subjected to developmental failure at the embryonic stage. Following this, we internally crossed the knockout mice and collected the embryo samples at the pre-implantation (E3.5 and E4.5), post-implantation (E7.5) as well as postnatal stage (Fig. 3a), and then observed the statistical loss of viable individuals per litter at the post-natal stage, decrement of normal gastrula as well as defective decidualization in the uterus at the early post-implantation stage (Fig. 3b,c and Extended Data Fig.4c). Meanwhile, the expression of mesoderm markers, *T* and *Mesp1*, in E7.5 GKO but normal morphological embryos seemed to show no obvious distinctions comparing to WT counterparts (Extended Data Fig. 4d). The comparable ratio of GKO embryo loss between post-implantation and postnatal stages illustrates the *in vivo* function of *Gm26793* may act earlier than the gastrulation stage. Additionally, the lack of aberrant mesoderm developmental phenotype in GKO embryos indicates that the *in vitro* mesodermal differentiation defect (Fig. 1g and Extended Data Fig. 2i,j) may be a byproduct of resistance to exit pluripotency and enhanced primitive endoderm differentiation capacity in GKO cells.

**Fig 3.**
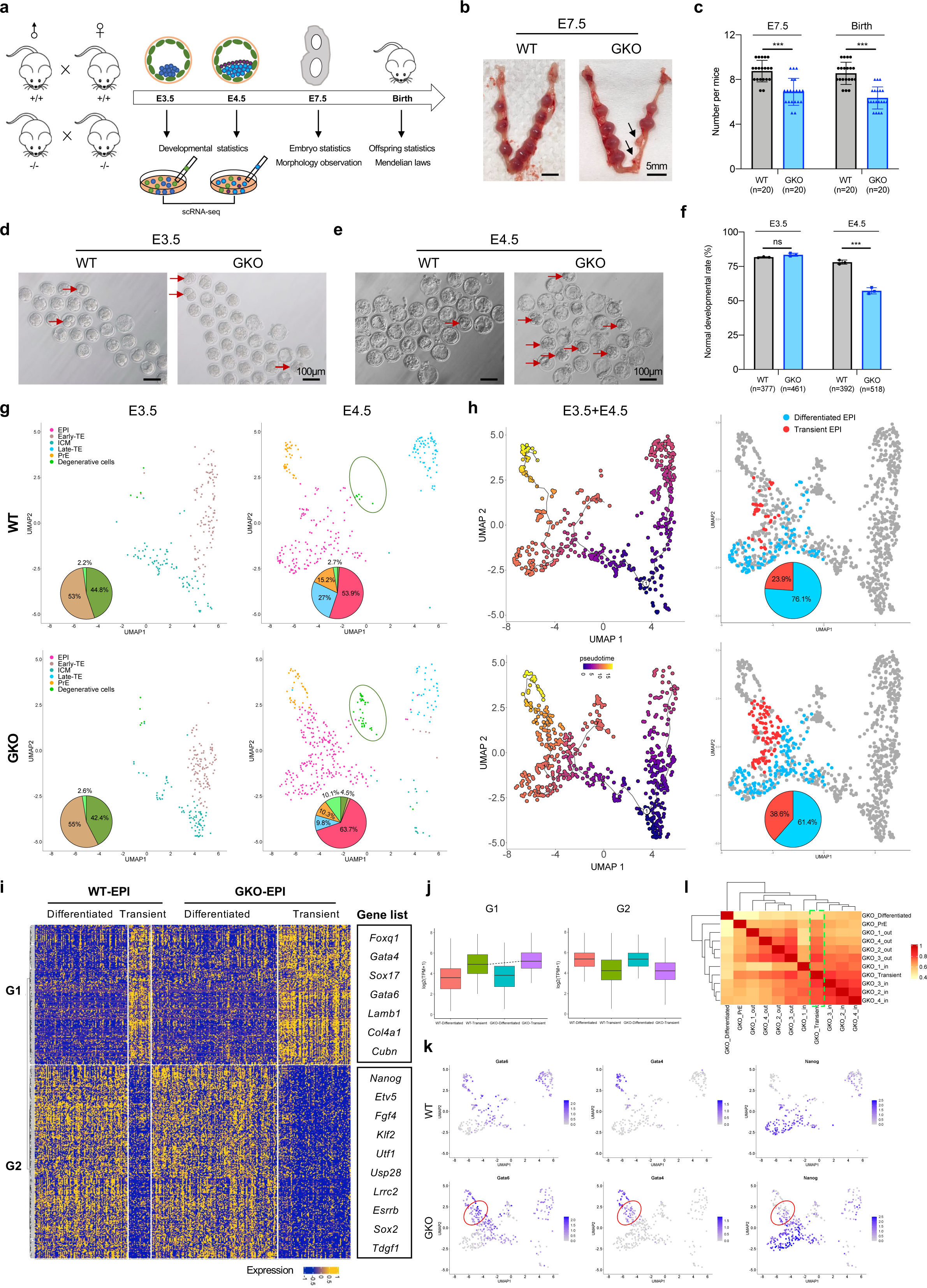
*Gm26793* null embryos exhibit developmental abnormalities during the lineage segregation between epiblast and primitive endoderm. (a) Experimental overview of mouse breeding, embryo preparation and the functional assays conducted at indicated developmental stages. (b) Representative images showing the resorbed decidua in GKO mouse at E7.5, which were highlighted by the arrows. Scale bar, 5 mm. (c) Quantification of the normal decidua collected at E7.5 and the acquired offspring number per litter. n represents the number of mating females. (d-e) Representative images showing the embryos collected at early (E3.5) (d) and late (E4.5) (e) blastocyst stage for both WT and GKO groups. Abnormal embryos were indicated by red arrows. Scale bar, 100 μm. (f) Quantification of the normal developmental rates at early and late blastocyst stage. N represents the total number of blastocysts collected. Data were summarized from three biologically independent experiments. (g) UMAP plots of scRNA-seq data from E3.5 and E4.5 blastocysts, respectively. The percentages of cell type composition were highlighted as pie chart embedded in related UMAP plot. Degenerative cells were circled by oval. (h) Pseudotime trajectories of pre-implantation cell lineages inferred by monocle 3 (left). Visualization of differentiated and transient epibast cells in UMAP plots based on the trajectory branch towards primitive endoderm (right). The percentages of epiblast subtype cells were highlighted as pie chart embedded in related UMAP plot. (i) Heatmap showing DEG groups among different epiblast subtypes displayed in (h). Representative genes were listed in the right panel. (j) Box plots showing average expression levels of differentiated and transient epiblast-specific genes in indicated cell clusters. (k) Antagonistic distribution of primitive endoderm and epiblast marker genes in transient epiblast cells. (l) Heatmap showing the transcriptomic correlation within *in vitro* EBs, differentiated and transient epiblast. All data were shown as means ± SEM. Student’s *t*-test, *** *p*<0.001.

To deeply delve into the potential function of *Gm26793* in the pre-implantation embryo, we collected early embryos at both E3.5 and E4.5 stages (Fig. 3d,e), in which the embryos start to form blastocyst consisting of three distinct lineages, trophoblast, epiblast, and primitive endoderm ^79^, and found the developmental rate of normal embryos between WT and GKO group seems to be equivalent at the E3.5 stage (Fig. 3d,f). However, once the embryos developed to the E4.5 stage, about 20% of GKO embryos exhibit blastocyst cavity formation defects (Fig. 3e,f). These results indicate that GKO embryos start to display developmental defects in the pre-implantation stage from E3.5 to E4.5.

Next, to systematically dissect the molecular and cellular changes of aberrant blastocysts caused by *Gm26793* knockout, we conducted single-cell RNA sequencing (scRNA-seq) of both WT and GKO embryos at E3.5 and E4.5 stage by using SMART-seq2 sequencing (Fig. 3a) ^80^. A total of 439 cells from WT embryos and 568 cells from GKO embryos were collected. Based on uniform manifold approximation and projection analyses (UMAP) and marker gene expression (Supplementary Table 4), five known cell clusters were identified in the embryo samples, and annotated as epiblast (EPI), inner cell mass (ICM), early trophectoderm (Eearly-TE), late trophectoderm (Late-TE) and primitive endoderm (PrE), respectively (Extended Data Fig. 4e,f,h). Notably, we found that one distinct cell group showing degenerative features, which manifests as high level of apoptotic related mitochondrial gene expression and majorly harbors cells in GKO embryos at the E4.5 stage (Fig. 3g and Extended Data Fig. 4e-g). Next, to capture the potential developmental phenotypes of GKO embryo, we reconstructed the pseudotime lineages for cell types in both WT and GKO embryos using Monocle 3 ^81^ (Fig. 3h). Following the trajectory of epiblast and primitive endoderm lineages segregation from inner cell mass, we determined two states of epiblast cells, transient and differentiated states, in both WT and GKO embryos (Fig. 3h). Statistic analysis identified that a greater proportion of GKO epiblast cells (38.6%) than WT epiblast cells (23.9%) are in transient state, which express higher level of primitive endodermal genes (G1) but lower level of pluripotent genes (G2) than epiblast in differentiated state (Fig. 3h-k and Extended Data Fig.4i, Supplementary Table 5). By integration with transcriptome data from *in vitro* EBs (Fig. 2c), we found that GKO_Transient cells also display high correlation with the inner and outer layer cells from GKO EBs (Fig. 3l). As known, the lineage segregation of ICM cells into epiblast and primitive endoderm relies on an intricate balance between pluripotent genes (such as *Nanog*) and primitive endoderm regulators (such as *Gata6*) ^82–84^. The dis-organized expression of epiblast genes or primitive endoderm genes in transient state epiblast cells caused by *Gm26793* knockout, similar with the *in vitro* stem cell differentiation defects, can disrupt the proper lineage segregation process and further induce defects of blastocyst development *in vivo*.

### *Gm26793*-mediated regulation is independent of the transcript and local transcriptional acitivity

To examine the roles of *Gm26793* transcriptional elongation, RNA molecules or its genomic locus in cell fate determination, we generated cells by deleting the promoter region upstream of transcriptional start site (TSS) of *Gm26793* (GPKO cells) (Fig. 4a and Extended Data Fig. 5a,b), or reintroduced full-length *Gm26793* transcripts into the GKO cells through lentivirus infection (GKO+*Gm26793*). Of note, the disruption of local transcription event by knocking-out of the promoter, which leads to the great reduction of *Gm26793* transcript (Fig. 4b,c), has limited effects on the differentiation towards mesoderm or primitive endoderm fate (Fig. 4d-f). Moreover, even though overexpression of *Gm26793* could sufficiently restore the expression of *Gm26793* in GKO cells during both EB and XEN differentiation (Fig. 4b,c), neither mesoderm markers nor primitive endoderm markers could be rescued (Fig. 4d-f).

**Fig 4.**
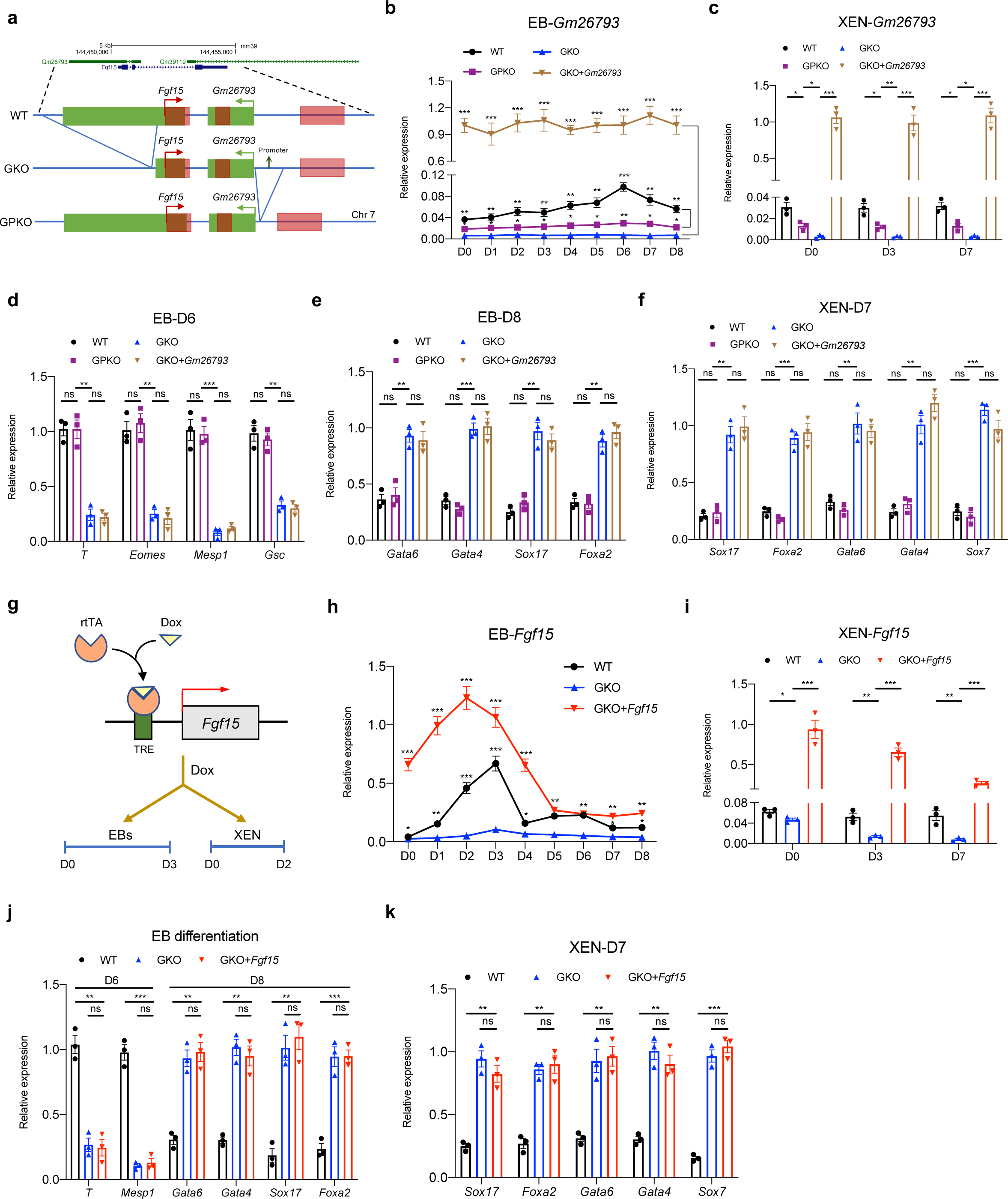
*Gm26793*-mediated regulation is independent of the transcript and local transcriptional activity. (a) The schematic diagram of the genomic information around *Gm26793* locus and related promoter knockout strategy. (b-c) Time-series expression of *Gm26793* in GPKO and GKO+*Gm26793* cells (*Gm26793* overexpression in GKO cells) during EB (b) and XEN (c) differentiation. (d-e) qPCR analyses of mesodermal (d) and PrE-related (e) genes on day 6 and day 8 of EB differentiation. (f) qPCR analyses of PrE-related genes in indicated cell groups on day 7 of XEN differentiation. (g) The workflow for overexpressing *Fgf15* during differentiation using tet-on inducible system. (h-i) The expression dynamics of *Fgf15* in indicated cell groups during EB (i) and XEN (j) differentiation. GKO+*Fgf15* represents conditional overexpression of *Fgf15* in GKO cells. (j-k) qPCR analyses of lineage-specific genes among respective cell types in EB (j) and XEN (k) differentiation. All qPCR data were shown as means ± SEM. Student’s *t*-test, \**p*<0.05, ** *p*<0.01, *** *p*<0.001.

It has been reported that the local transcriptional activity could affect the expression of both lncRNA and its nearby protein-coding gene ^85, 86^. In this study, *Gm26793* is located at the divergent direction of the *Fgf15* locus with partial genomic overlap (Fig. 4a), and the knockout of *Gm26793* indeed leads to the down-regulation of *Fgf15* expression during both EB and XEN differentiation (Fig. 4h,i). To explore the role of *Fgf15*, we conditionally over-expressed *Fgf15* in GKO cells (named as GKO+*Fgf15*) and then performed stem cell differentiation (Fig. 4g-i). However, inducible over-expression of *Fgf15* at the early stage of differentiation has no effects on rescuing mesendoderm and primitive endoderm defects caused by *Gm26793* genomic knockout (Fig. 4j,k). Collectively, these results support that the transcriptional activity and transcripts of *Gm26793*, as well as adjacent coding gene, are dispensable for primitive endoderm differentiation.

### *Gm26793* regulates stem cell differentiation through direct inter-chromosomal interaction with *Cubn* locus

Next, we want to test whether the genomic locus of *Gm26793* could contribute to the differentiation defects, especially through remote chromatin interactions. To this end, we performed circular chromosome conformation capture sequencing (4C-seq) using the *Gm26793* knockout region as a bait (Fig. 5a) to query genome-wide chromatin interactions. Among 1919 interacting targets, 1318 nearest neighboring genes (Supplementary Table 7) were finally assigned in three replicates of WT mESCs (Fig. 5b). Besides the relatively higher enrichment of intra-chromosomal peaks, we also detect the pervasive existence of interacting chromatin hits at the neighboring chromosomes (Fig. 5b). These results denote that the genomic locus of *Gm26793* may act as a chromatin interaction hub by forming both intra- and inter-chromosomal interactions.

**Fig 5.**
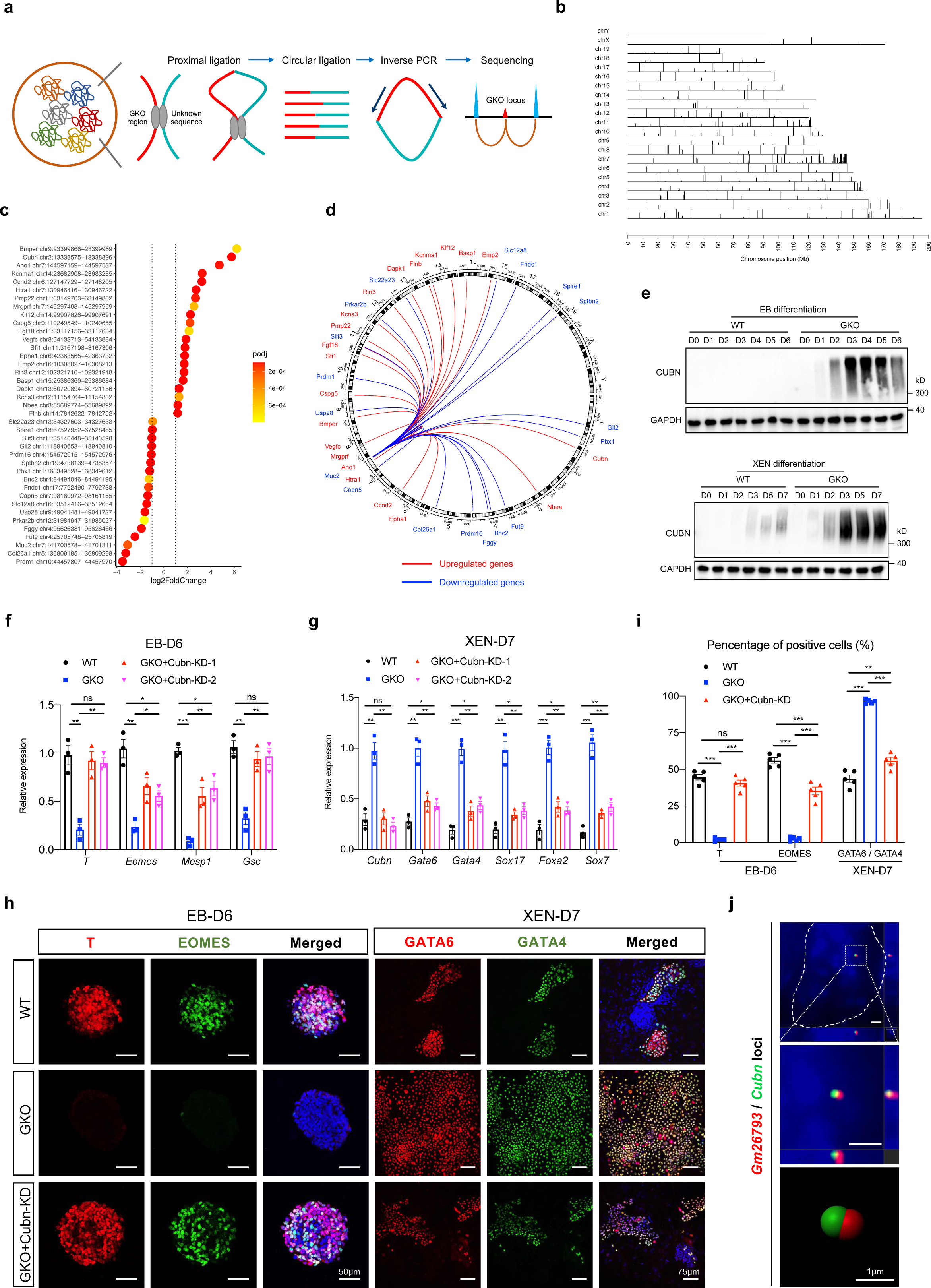
The genomic locus of *Gm26793* forms inter-chromosomal interaction with *Cubn*. (a) Schematic representation describing the basic workflow for the 4C assay. The cyan line indicates potential sequences interacting with *Gm26793* locus. (b) The genome-wide distribution of *Gm26793* interacting peaks captured by 4C-seq. (c) Dot plot summarizing genes with both significant *Gm26793* interacting hits revealed by 4C-seq and obvious expression changes between WT and GKO cells based on transcriptomic data. Genes were ranked by the fold change of expression. (d) Circos plot showing inter- and intra-chromosomal connectivity associated to the genomic region of *Gm26793*. Line colors represent up-(red) and down-regulation (blue) of annotated genes. (e) Western blotting results reporting the aberrant activation of CUBN in GKO cells during EB (top) and XEN (bottom) differentiation. (f-g) The expression recovery of mesoderm-associated genes on day 6 of EB differentiation (f) and PrE-related genes on day 7 of XEN differentiation (g) upon *Cubn* knockdown in GKO cells. (h-i) Representative images describing the rescue of protein distribution during EB (left) and XEN (right) differentiation. Scale bar, 50 μm and 75 μm. Quantification of the data was shown in (i). (j) dCas9-mediated living cell imaging showing the co-localization of *Gm26793* (Red probe) and *Cubn* (Green probe) loci in WT mESCs. Scale bar, 1 μm. All data were shown as means ± SEM. Student’s *t* test, \**p*<0.05, ** *p*<0.01, *** *p*<0.001.

To refine the specific functional targets for *Gm26793* locus, we combined the transcriptome data (Supplementary Table 6) of WT and GKO mESCs with 4C-seq data, and screened out 39 gene candidates with significant gene expression changes (21 genes were up-regulated, 18 genes were down-regulated) triggered by *Gm26793* knockout (Fig. 5c, d and Extended Data Fig. 5c,d). Gene ontology analysis revealed that these genes were highly involved in epithelium development (Extended Data Fig. 5e). RT-qPCR analyses further corroborated that 18 out of 21 up-regulated genes and 13 out of 18 down-regulated genes were truly altered in GKO cells (Extended Data Fig. 5f,g). Furthermore, by referencing the published datasets ^87–92^, we eventually focused on eight genes with potential roles in stem cell differentiation, including five upregulated genes (*Cubn, Ano1*, *Htra1*, *Sfi1*, and *Flnb*) and three down-regulated genes (*Slc12a8*, *Usp28*, *Fut9*). Then, we established cell lines with knocking-down the up-regulated genes or over-expressing the down-regulated genes in GKO cells, respectively (Extended Data Fig. 5h). The restoration of 7 selected genes (*Ano1*, *Htra1*, *Sfi1*, *Flnb*, *Slc12a8*, *Usp28*, *Fut9*) failed to recover the expected cell lineage transition during EBs and XEN differentiation (Extended Data Fig. 5i,j). On the contrary, for the gene of *Cubn*, which was up-regulated in GKO cells during differentiation and linked by inter-chromosomal interaction with *Gm26793* locus (Fig. 5c-e and Extended Data Fig. 6a-c,h), we found that knockdown of *Cubn* expression level in GKO cells could partially rescue the differentiation phenotype (Fig. 5f-i and Extended Data Fig. 6c-g). These results indicate that *Cubn* could be the direct interaction target of *Gm26793*, and the elevated expression of *Cubn* should be responsible for the raised primitive endoderm differentiation potential in GKO cells. Subsequently, to unambiguously verify the inter-chromosomal association between *Gm26793* and *Cubn* loci, we took advantage of the CRISPR-dCas9 assisted live imaging system ^93^ and designed specific crRNA probes targeting the genomic locations of *Gm26793* and *Cubn*, to visualize the spatial distribution of each locus in the WT living mESCs. As predicted, the genomic loci of *Gm26793* and *Cubn* are spatially co-localized in the nuclei (Fig. 5j).

Taken together, these data unveil that *Gm26793* in chromosome 7 actually forms inter-chromosomal interaction with *Cubn* in chromosome 2, and the specific deletion of *Gm26793* locus may release this contact and enhance the expression level of *Cubn* during stem cell differentiation process, which supports the notion that the inter-chromosomal organization between *Gm26793* and *Cubn* loci can behave as a molecular lock to restrict the expression of *Cubn* in WT cells.

### The pervasive remodeling of epigenomic landscape in GKO cells could be restored by silencing *Cubn* expression

To understand the molecular basis of the facilitated primitive endoderm differentiation potential decribed above, we analyzed the global epigenomic pattern in both WT and GKO cells by profiling the genomic distribution of chromatin-accessible regions, active histone marker-H3K27ac, as well as promoter-related histone marker-H3K4me3 (Fig. 6a) and observed a consistent elevation of chromatin accessibility, H3K4me3 and H3K27ac enrichment around *Cubn* locus (Fig. 6b). By systematic comparison between WT and GKO cells, we found that the epigenomic pattern including chromatin accessibility and histone modification distributions were pervasively altered, in which the global distribution of H3K27ac exhibits the most dramatic changes in GKO cells (Fig. 6c and Extended Data Fig. 7a, Supplementary Table 8). Considering that the enrichment of H3K27ac has been used as an epigenetic marker to demarcate the primed and activated chromatin states of regulatory elements, the dramatic alteration of H3K27ac in GKO cells indicates that widespread chromatin state transition may occur in GKO cells, which may be related to the enhanced responsiveness to primitive endoderm. Following this, we focused on analyzing chromatin regions with up-regulated H3K27ac in GKO cells and further clustering these regions based on the existence of promoter-related histone marker-H3K4me3 (Fig. 6d). By dividing these regions into newly derived active promoters (H3K4me3 and H3K27ac double positive), and newly derived active enhancers (H3K27ac positive only) in GKO cells, we found that these newly derived regulatory elements were mostly enriched around endoderm-related genes, such as *Gata6*, *Gata4*, *Sox17* and *Sox7* (Fig. 6e-g and Extended Data Fig. 7b,c, Supplementary Table 9). Besides, we found that the relative enrichment of H3K27ac at both active enhancers and promoters in GKO cells can be largely erased by knocking down *Cubn* expression, and return to a comparable level with WT cells (Fig. 6e,g and Extended Data Fig. 7d). Thus, the genetic knockout of *Gm26793* leads to pervasive remodeling of the epigenomic landscape, especially for global re-distribution of active markers-H3K27ac around endodermal genes, and the remodeling of active H3K27ac modification can be largely revised by knocking-down *Cubn* expression.

**Fig 6.**
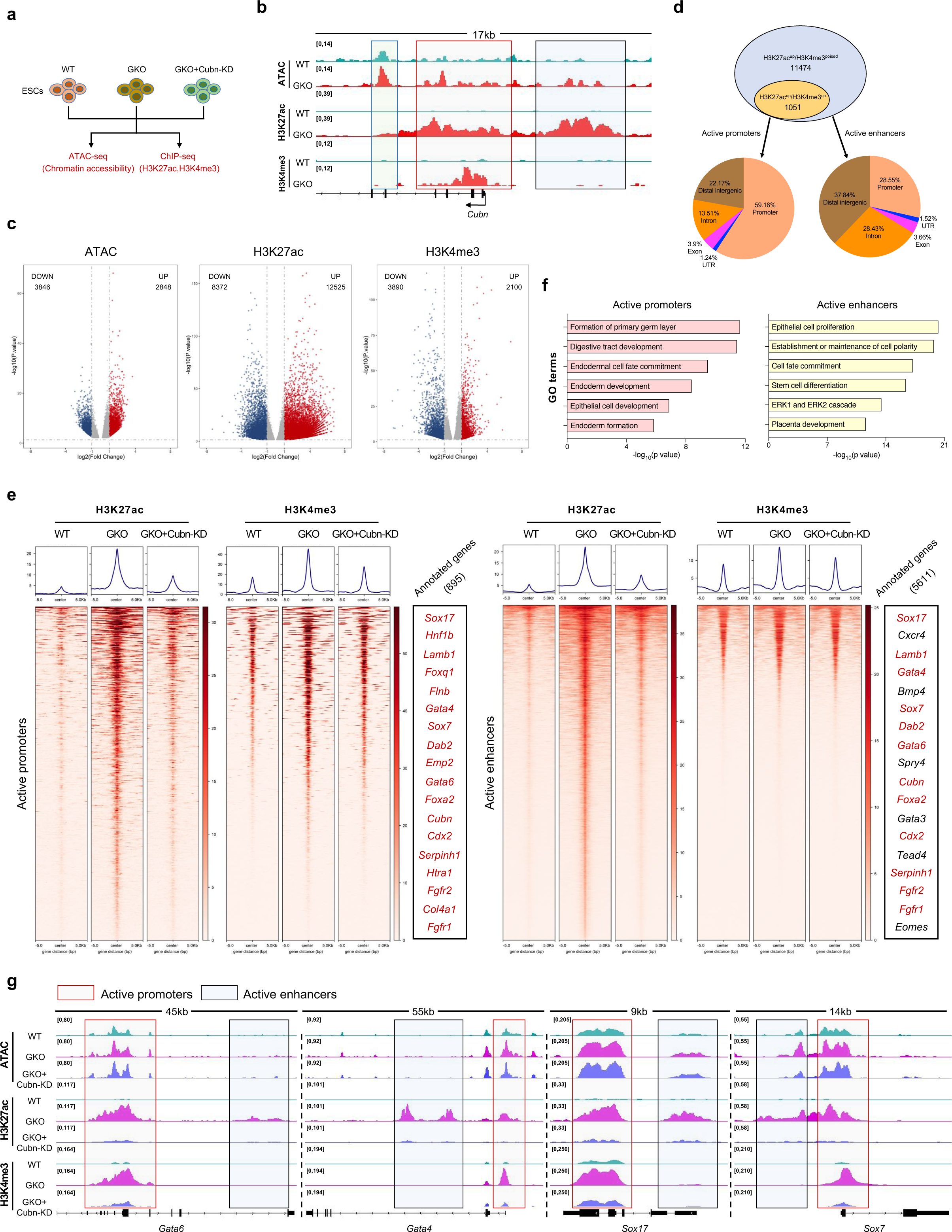
*Gm26793* knockout leads to pervasive epigenomic remodeling, which can be rescued by knocking-down *Cubn* expression. (a) Schematic of epigenomic profiling among related mESCs. (b) Genome browser snapshot showing the increased chromatin accessibility, H3K27ac and H3K4me3 enrichment around *Cubn* locus in GKO mESCs. (c) Volcano plot determining the differential distribution of chromatin accessibility (left), H3K27ac (middle), and H3K4me3 (right) between WT and GKO cells. Enhanced signals in GKO cells were highlighted in red, decreased signals in GKO cells were highlighted in blue. (d) Venn diagram showing genomic regions with increased H3K27ac in GKO cells, which were analyzed and divided into active promoters and active enhancers based on the existence of H3K4me3 modification. Genomic annotations of divided regions were presented as pie chart. (e) Heatmaps showing the signal enrichment of H3K27ac and H3K4me3 around selected genomic regions (active promoters: left; active enhancers: right). Representative annotated genes for the selected regions were also shown. Identical genes are labelled in red. (f) GO term enrichment for the genes in active promoter group (left) and active enhancer group (right). (g) Representative genome browser snapshot showing efficient recovery of epigenetic distribution in GKO+Cubn KD cells. Promoter regions were highlighted in light red rectangles, enhancer regions were highlighted in light blue rectangles.

### CTCF and cohesin complex acts as a lock ring to fix the inter-chromosomal interaction between *Gm26793* and *Cubn* loci

Previously, ChIA-PET analyses showed that CTCF and cohesin extensively participate in the inter- and intra-chromosomal interactions and configure the genome into distinct domains that possess unique epigenetic states ^26, 44, 94^. To further delineate the mechanism of the inter-chromosomal molecular lock within *Gm26793*-*Cubn* loci, we performed CTCF and RAD21 ChIP-seq in both WT and GKO mESCs. Globally, the genomic distributions of CTCF and RAD21 within peak center ± 5 kb in both cells remain largely unchanged (Extended Data Fig. 8a and Supplementary Table 10). As to *Gm26793* and *Cubn* loci, we found both CTCF and RAD21 are co-bound at the *Gm26793* knockout region, as well as around the *Cubn* locus (Fig. 7a). These results infer that the inter-chromosomal lock between *Gm26793* and *Cubn* may be fixed by the co-binding of CTCF-cohesin complex, just like a lock ring, at the highlighted regions.

**Fig 7.**
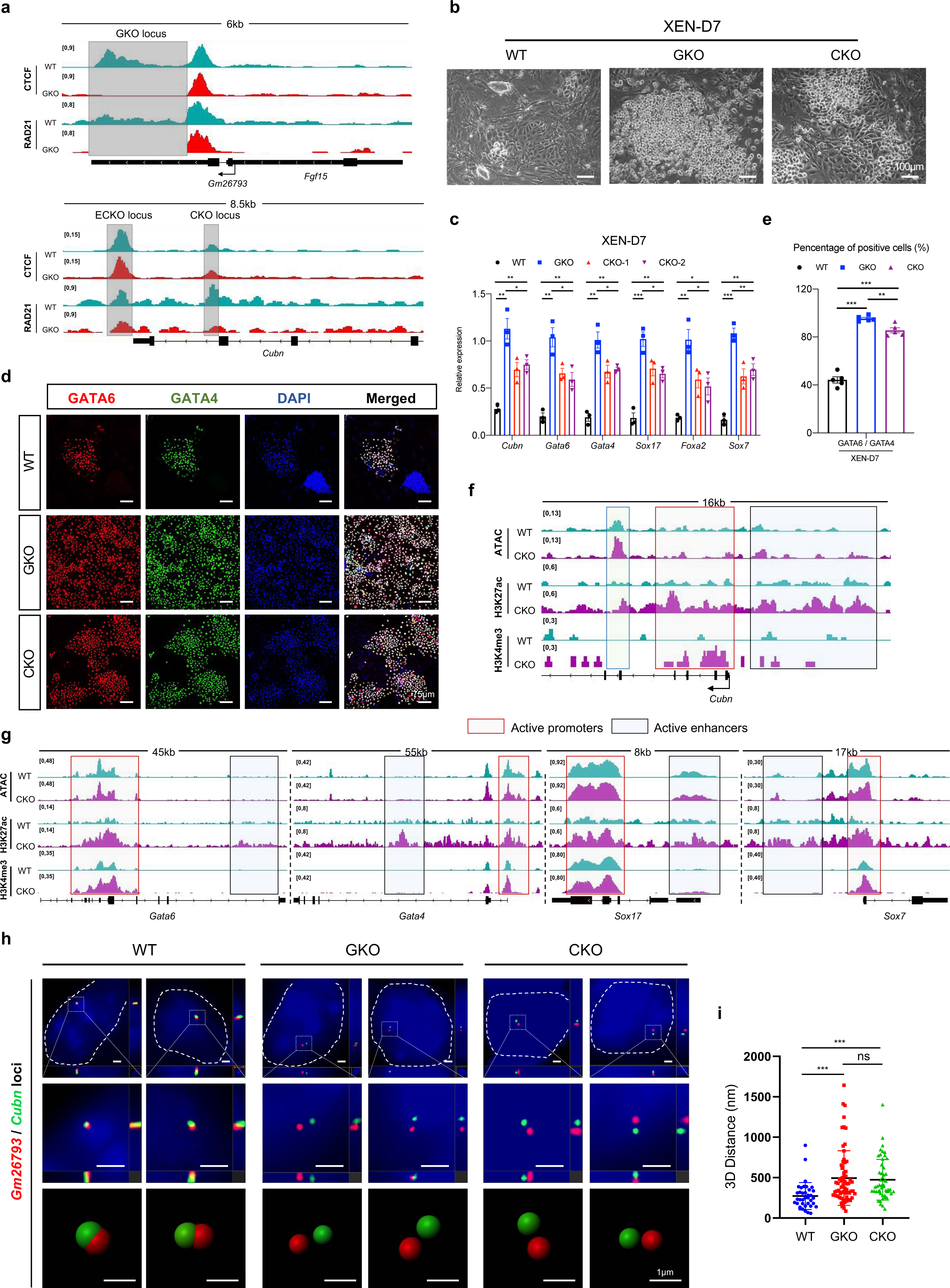
The binding of CTCF-cohesin complex orchestrates the inter-chromosomal association between *Gm26793* and *Cubn* loci. (a) Genome browser view demonstrating the binding of CTCF and cohesin at *Gm26793* and *Cubn* loci in WT and GKO mESCs. Grey boxes represent the co-binding sites of CTCF and RAD21, which were subjected to genomic knockout in GKO cells and following experiments. (b-c) Morphologies of differentiated cells at bright field (b) and relative expression level of PrE-related genes (c) in each cell type as indicated. Scale bar, 100 μm. (d-e) Representative images displaying the expression of GATA6 and GATA4 protein in XEN differentiated from WT, GKO and CKO mESCs. Statistical analyses of fluorescent signal were summarized in (e) as bar plot. Scale bar, 75 μm. (f-g) Genome browser view showing the dynamics of chromatin accessibility, H3K27ac and H3K4me3 signals around *Cubn* locus (f) and PrE-related gene regions (g) in WT and CKO mESCs. (h-i) Live cell imaging and 3D-reconstruction recording the spatial distribution of *Gm26793* (Red) and *Cubn* (Green) loci in indicated mESCs. Scale bar, 1 μm. The 3D-distances between *Gm26793* and *Cub*n loci were presented in (i) (WT, n=37 cells; GKO, n=65 cells; CKO, n=51 cells). All data were shown as means ± SEM. Student’s *t* test, \**p*<0.05, ** *p*<0.01, *** *p*<0.001.

To assess the roles of CTCF-cohesin complex enrichment, we generated two ESC lines with CTCF-cohesin binding sites depletion within *Cubn* non-coding region (named as ECKO and CKO, respectively) (Extended Data Fig. 8b,c,e,f), and performed XEN differentiation afterwards. For the ECKO cells, we found that the expression of *Cubn*, as well as the primitive endoderm markers, has not been changed (Extended Data Fig. 8d). In contrast, the expression of *Cubn* was evidently up-regulated in CKO cells, albeit with a less extent than GKO cells (Extended Data Fig. 8g). Further differentiation towards the primitive endoderm fate of CKO cells also showed the appearance of boosted primitive endoderm differentiation preference, which recapitulated the differentiation phenotype of GKO cells (Fig. 7b-e). Moreover, epigenomic profiling also showed the increment of active epigenomic features at endoderm-related genes, including *Cubn*, *Gata6*, *Gata4*, and *Sox17,* in CKO cells (Fig. 7f,g and Extended Data Fig. 7d). Next, to faithfully reflect the indispensability of CTCF-cohesin complex binding in orchestrating the *Gm26793*-*Cubn* inter-chromosomal association, we applied the CRISPR-dCas9 assisted imaging technology to visualize the spatial localization of both loci in WT and anchor sites deletion cells (GKO and CKO cells) and discovered the originally tight co-localized *Gm26793* and *Cubn* loci were separated apart in both GKO and CKO cells (Fig. 7h and Extended Data Fig. 8h). Moreover, statistical measurement of the spatial distance between *Gm26793* and *Cubn* revealed that the 3D distance reached to a comparable level in GKO and CKO cells, which highlighted the equivalent roles of both *Gm26793* and CKO loci in forming the inter-chromosomal interaction (Fig. 7i). These results indicate that the binding of CTCF-cohesin complex at GKO and CKO regions is directly responsible for establishing the inter-chromosomal contact between *Gm26793* and *Cubn* loci.

## Discussion

In this study, we identify a germ layer-specific lncRNA gene, *Gm26793,* through comprehensive analyses of the spatial transcriptome atlas of the mouse gastrula (Fig. 1a). To investigate its functional significance, we establish GKO mESCs and mouse models by genetically knocking-out *Gm26793*. During *in vitro* spontaneous differentiation, we find a significant impairment in the expression of mesendodermal and mesodermal genes (Fig. 1e and Extended Data Fig. 3e), along with abnormal up-regulation of genes related to primitive endoderm development (Fig. 2a and Extended Data Fig. 3a). Meanwhile, about 20% of embryos lacking *Gm26793* locus also exhibit developmental arrest during the lineage segregation between epiblast and primitive endoderm fate, which further results in decidualization abnormalities at the post-implantation stage, as well as decrement of normal individual per litter during the mouse life cycle (Fig. 3b-f). Towards an in-depth understanding of the molecular basis for *Gm26793* function, we re-express *Gm26793* and the neighboring *Fgf15* gene in GKO cells, respectively. However, no functional rescue could be observed, indicating the function of *Gm26793* is independent of its own transcript and proximal gene (Fig. 4). As revealed by 4C-seq and live cell FISH, we unravel the genomic locus of *Gm26793* in chromosome 7 can directly interact with *Cubn* (Fig. 5d,j), a primitive endoderm regulator located in adjacent chromosome 2. Molecularly, this specific inter-chromosomal interaction is bridged by CTCF-cohesin complex (Fig. 7a), and genomic depletion of CTCF-cohesin anchor sites residing in *Gm26793* or *Cubn* locus equally breaks up the inter-chromosomal interaction, releases the restriction of *Cubn* expression, remodels the epigenomic landscape in mESCs, and finally forms a primitive endoderm signal-responsiveness state (Fig. 6 and 7). Notably, these effects can be restored by knocking-down of *Cubn* (Fig. 5f-i and 6e). Taken together, we propose an ‘inter-chromosomal molecular lock-ring’ model to clearly elucidate the necessity of *Gm26793* locus in controlling *Cubn* expression and directing mouse blastocyst development (Fig. 8).

**Fig 8.**
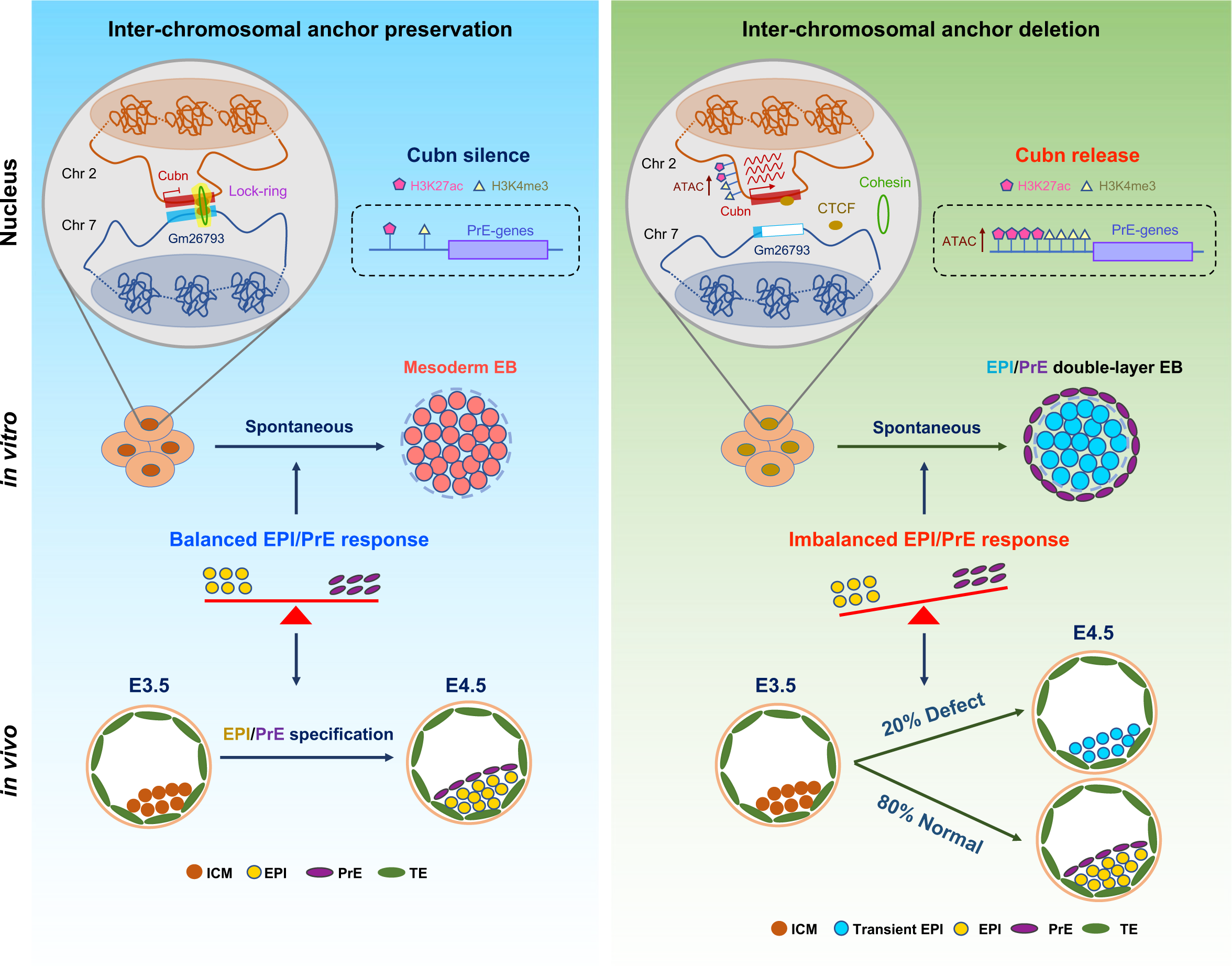
The molecular lock-ring model highlighting the function and mechanism of *Gm26793*-*Cubn* inter-chromosomal interaction in PrE lineage commitment.

We determine that *Gm26793* functions as a crucial regulator for both stem cell differentiation and early embryogenesis. lncRNAs have been revealed to fine-tune biological processes and regulate the spatiotemporal expression of pleiotropic developmental loci instead of being master regulators or switches of development ^95^. A considerable portion of lncRNAs has been reported to be dispensable, albeit with context-dependent function during mammalian development ^96^. Thus, a comprehensive evaluation of the functional relevance of lncRNA genes warrants dual integration of *in vitro* and *in vivo* systems. Here, we generate both embryonic stem cell and mouse model with *Gm26793* locus removal. As revealed in the *in vitro* spontaneous differentiation system, the GKO EBs, which are supposed to express mesodermal genes, exhibit two-layer structures with primitive endodermal genes enriched at the outer layer and pluripotent genes enriched at the inner layer (Fig. 2d). Hence, for the *in vitro* system, *Gm26793* knockout either leads to cell arrest at the pluripotent state, or those, which successfully exit the pluripotent state, prone to differentiate into primitive endoderm fate. Consistently, in the *in vivo* mouse model, we detect a substantial incidence of blastocyst formation defects in GKO embryos (Fig. 3e,f). As revealed by single cell RNA-seq results of relevant embryos, we observe the over-representation of epiblast cells in degenerative or transient states for GKO embryos (Fig. 3g,h), which underscores the reason for higher frequency of developmental failure upon *Gm26793* deficiency. The formation of normal blastocyst relies on timely separation and balanced expression of lineage-specific genes between epiblast and primitive endoderm cells ^97–99^. The elevated porportion of transient epiblast cells, which act as the *in vivo* counterpart for the *in vitro* GKO EBs with strong co-expression of pluripotent and primitive endodermal genes (Fig. 3i-l), disrupts of the celluar homeostasis and sequential gene expression programs for the lineage segregation during early embryogenesis.

We report the existence of functional inter-chromosomal interaction involving lncRNA gene. As a newly defined regulatory dimension, various lncRNAs have been identified and reported to be involved in the modulation of chromatin function, alteration of cytoplasmic mRNAs’ stability and translation, as well as interference of signaling pathways ^100, 101^. In many cases, the abundance of lncRNA transcript and the adjacent genes have been implicated in their biological functions. However, in this study, the restoration of *Gm26793* or the neighboring *Fgf15* gene fails to rescue the developmental defects caused by *Gm26793* knockout (Fig. 4). Investigation of the correspondent genomic locus by capturing potential interacting chromatin regions reveals the existence of direct inter-chromosomal interaction between *Gm26793* and *Cubn* loci (Fig. 5j). The formation of a direct chromatin-chromatin interacting loop has been treated as a common mechanism employed by genomic regulatory elements, such as enhancers, promoters. Genomic elements with enhancer activities are usually involved in the maintenance or up-regulation of target gene expression. However, in our study, although direct chromatin interaction can be identified between *Gm26793* and *Cubn*, *Gm26793* seems to be a repressive molecular lock, which can silence *Cubn* expression in normal mESCs (Fig. 5c). This suggests that *Gm26793* may operate through a mechanism similar to recently identified regulatory elements known as silencers ^102^. A more comprehensive characterization of epigenetic features and sequence composition of *Gm26793* will facilitate the understanding of how this molecular lock works.

We identify CTCF-cohesin complex as the crucial architectural mediator that establishes the inter-chromosomal lock between *Gm26793* and *Cubn*. As shown in Fig. 7, we find that both CTCF and cohesin can specifically bind to the genomic locus of *Gm26793* and *Cubn*. Genetic removal of individual CTCF coupling anchor site (GKO or CKO locus) significantly disrupt the inter-chromosomal interaction, further resulting in the boosted responsiveness to primitive endoderm differentiation signals (Fig. 7c,h). As known, CTCF-cohesin complex has been recognized as one of the fundamental players in the “loop extrusion” model within the same chromosome, especially for the establishment of TADs ^103^. Since the specific interaction occurs between two distinct chromosomes in this study, it remains to determine whether CTCF-cohesin complex behaves similarly as in intra-chromosomal interaction, in composing the “inter-chromosomal genomic love story of kissing” ^104^. As revealed by the epigenomic profiling of H3K27ac, H3K4me3, and chromatin accessibility, a single knock-out of *Gm26793* or CKO locus leads to global pervasive remodeling of the epigenetic landscape (Fig. 6b,g and 7f,g). Accompanying complex molecular cascades possibly participate in response to the genetic alteration. Thus, we hypothesize that additional factors, like epigenetic factors, may exist in concert with CTCF-cohesin complex in manipulating this inter-chromosomal interaction. Further investigation of the epigenetic features and motif enrichment will provide valuable insights into the precise mechanisms of this molecular lock.

In conclusion, our observation extends the classical paradigm of how transcriptional regulation occurs through lncRNA, and demonstrates the existence and biological significance of inter-chromosomal interaction during stem cell differentiation and embryo development. Future studies related to molecular dynamics reflecting the formation and maintenance of inter-chromosomal interactions and subsequent molecular cascades will broaden the horizon of stem cell fate determination and mammalian embryogenesis.

## Extended Figure Titles and Legends

**Extended Data Fig 1.**
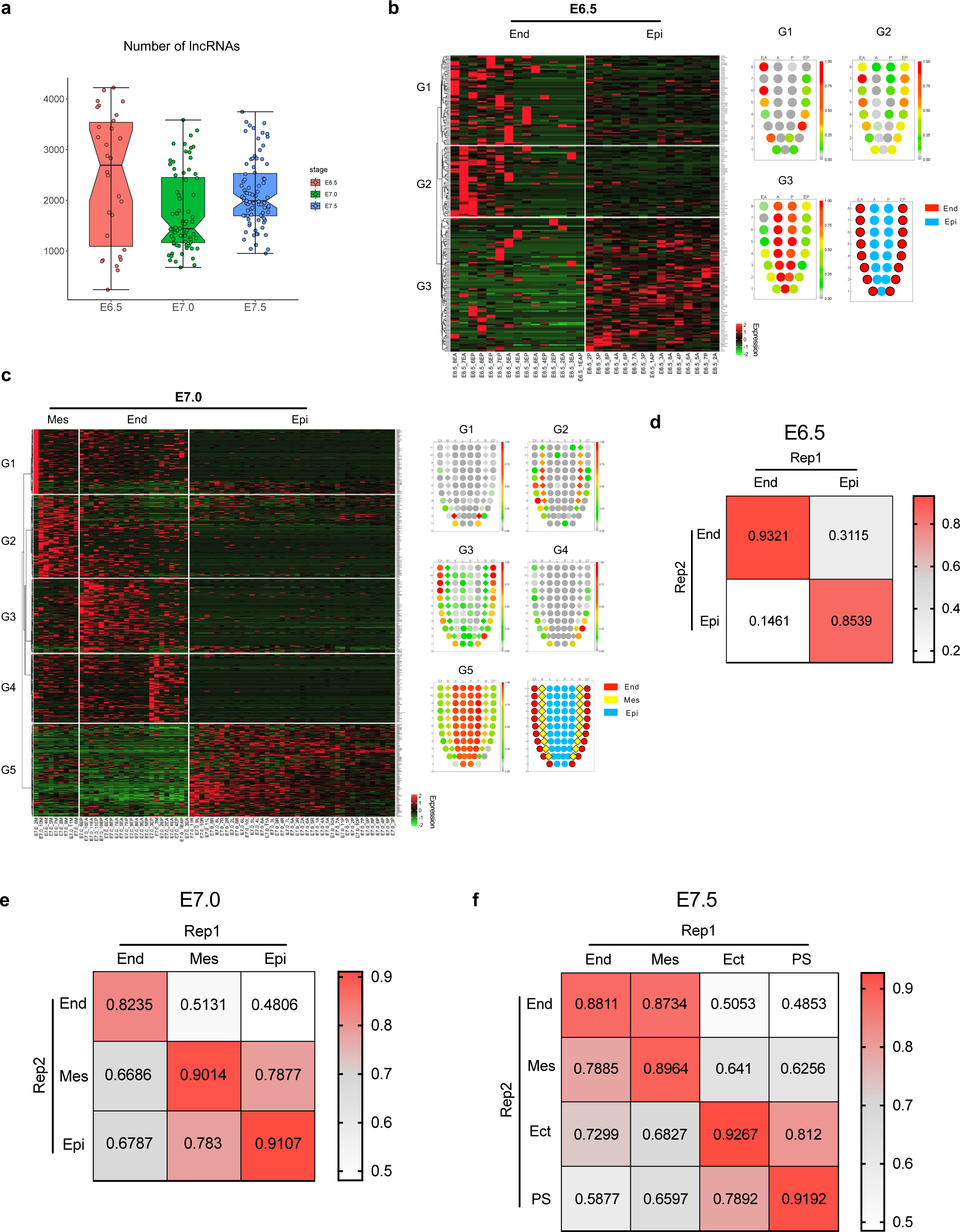
Differentially expressed lncRNAs during mouse gastrulation. (a) Box plot showing the number of detected lncRNAs for each GEO-seq samples from E6.5 to E7.5 mouse embryos. (b-c) Heatmaps and corn plots of lncRNAs with germ layer specific expression in E6.5 (b) and E7.0 (c) gastrula. (d-f) The spearman correlation coefficient of distinct germ layers between embryonic replicates at E6.5 (d), E7.0 (e) and E7.5 (f) stage based on the identified differentially expressed lncRNAs.

**Extended Data Fig 2.**
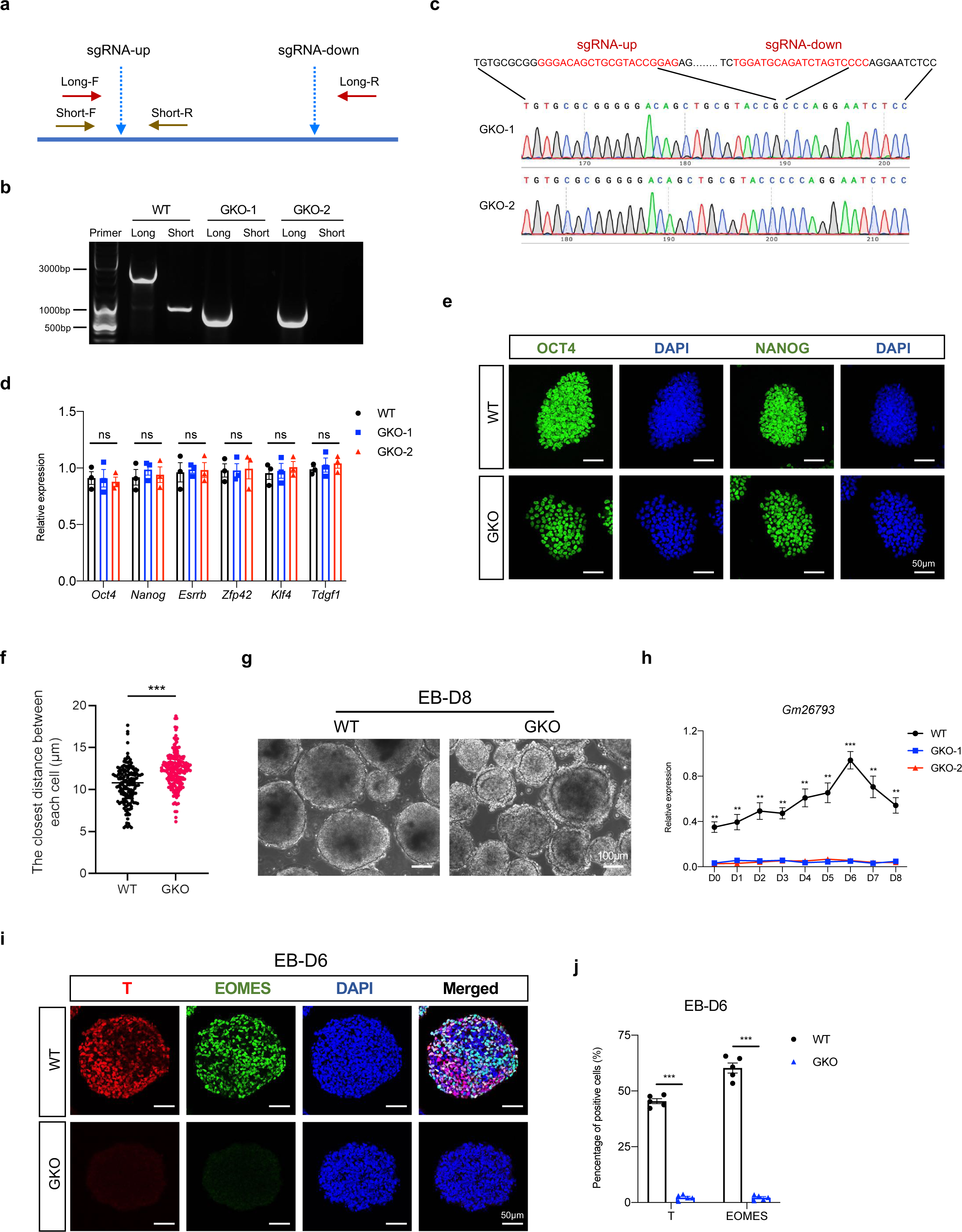
The establishment of GKO mESCs and expression identification of mesoderm-related genes during EB differentiation. (a) Schematic description of genotyping strategy and related primer designed in determining knockout clones. (b) Representative image of gel electrophoresis showing the genotyping PCR results of GKO mESCs. (c) Sanger sequencing results of the acquired GKO cells demonstrating successful removal of targeted allele in *Gm26793* locus. (d) Bar plot showing the comparable expression level of core pluripotency marker genes in WT and GKO mESCs. (e) Immunostaining analyses of OCT4 and NANOG in WT and GKO mESCs. Scale bar, 50 μm. (f) Statistic calculation of the spatial distance between each nucleus in WT and GKO mESCs. (g) The morphologies of differentiated EBs at day 8 in both WT and GKO group. Scale bar, 100 μm. (h) qPCR analyses determining the relative expression changes of *Gm26793* transcript during spontaneous differentiation. (i-j) Immunoflorescence analyses showing the distribution of T and EOMES protein in WT and GKO EBs collected at day 6. Scale bar, 50 μm. Quantification of the data was shown in (j) (n=5), n represents distinct EBs. All data were shown as means ± SEM. Student’s *t*-test, * *p* < 0.05, ** *p* < 0.01, *** *p*<0.001.

**Extended Data Fig 3.**
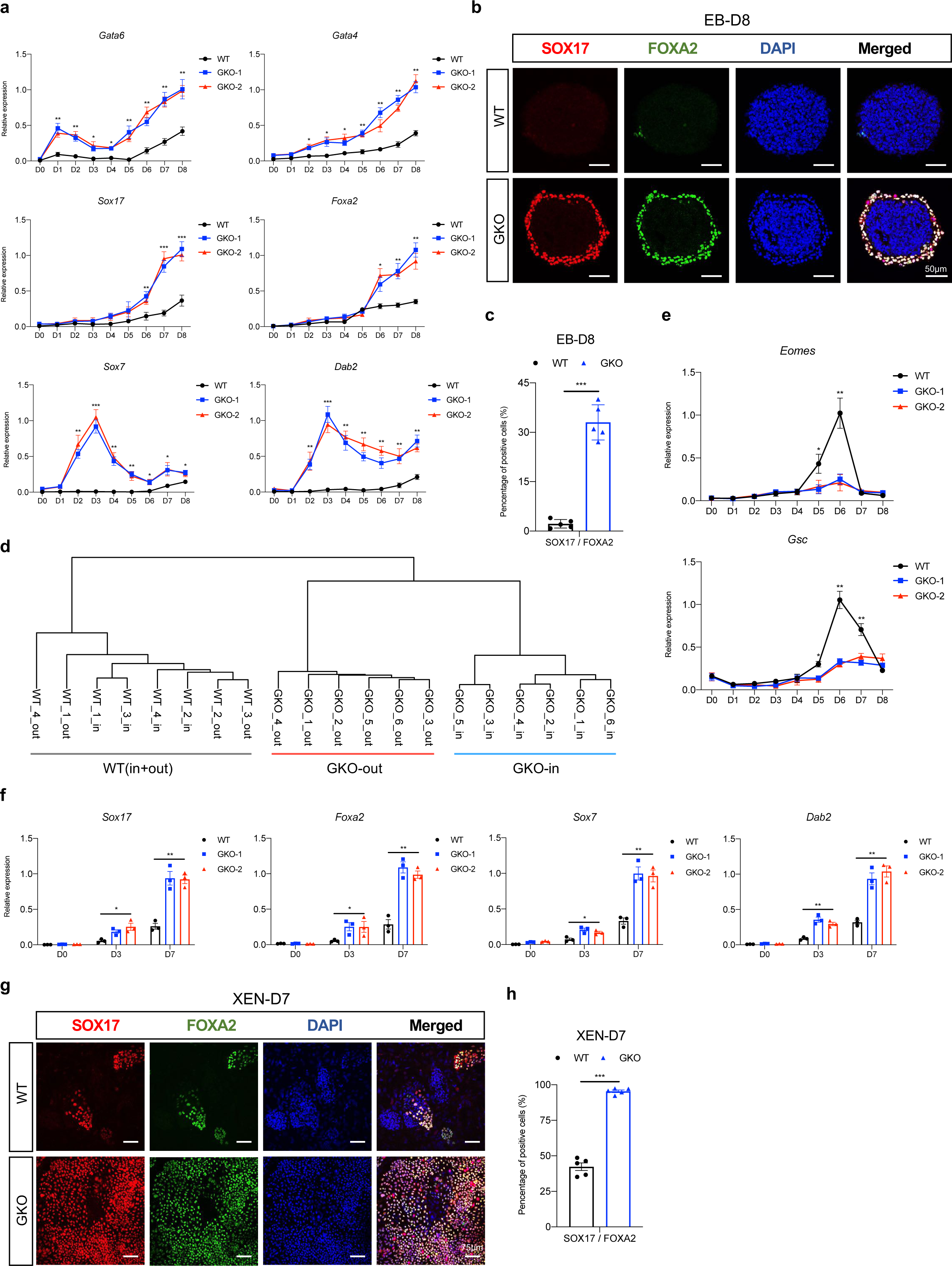
The expression of PrE marker genes were largely enhanced in GKO cells during EB and XEN differentiation. (a) qPCR analyses identify the immediate activation of PrE-related genes during EB differentiation upon *Gm26793* knockout. (b-c) Immunoflorescence analyses of SOX17/FOXA2 in WT and GKO EBs at day 8. Scale bar, 50 μm. Quantification of the data was shown in (C). (d) Hierarchical clustering based on the transcriptomic data of outside and inside EB differentiation samples. (e) The expression dynamics of mesendodermal markers-*Eomes* and *Gsc* during WT and GKO EB differentiation. (f) Significant upregulation of PrE-related genes during XEN differentiation upon *Gm26793* knockout. (g-h) Immunoflorescence analyses of SOX17/FOXA2 in WT and GKO XEN at day 7. Scale bar, 75 μm. Quantification of the data was shown in (h). All data were shown as means ± SEM. Student’s *t*-test, \**p*<0.05, ** *p*<0.01, *** *p*<0.001.

**Extended Data Fig 4.**
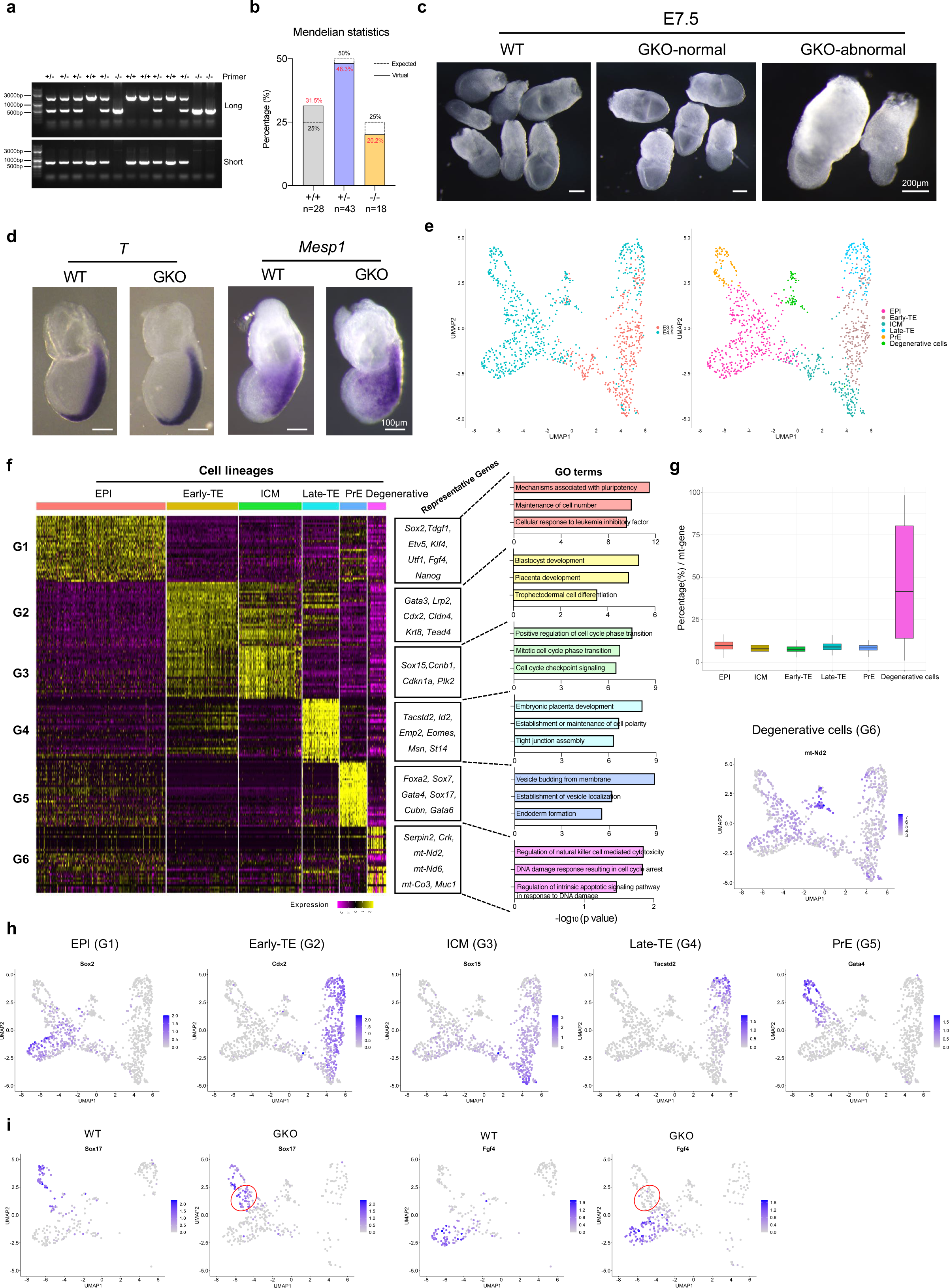
The characterization of GKO embryos and cell type identification of scRNA-seq. (a) DNA electrophoresis image showing the PCR genotyping of offspring born form GKO heterozygous mouse. (b) Bar plot depicting the mendelian statistics of mouse progeny acquired through crossing GKO heterozygous mouse. (c) Gross morphologies of WT embryos, GKO embryos with normal morphology, GKO embryos with abnormal morphology. (d) Whole-mount in situ hybridization showing the indistinguishable distribution of *T* and *Mesp1* in WT and GKO embryos collected at E7.5 stage. (e) UMAP plots of the gene expression profiles of individual cells regarding developmental stages (left) and cell types(right). The annotation for each cell cluster was included at the right side. (f) Heatmap showing the expression of cell type-specific signature genes based on the single-cell transcriptome atlas. Representative genes and GO terms of each DEG group were listed in the right panel. (g) The percentage of expressed mitochondrial genes in distinct cell populations. (h) Expression profile of selected lineage-specific genes in UMAP plots. (i) Expression pattern of *Sox17* and *Fgf4* in transient epiblast cells.

**Extended Data Fig 5.**
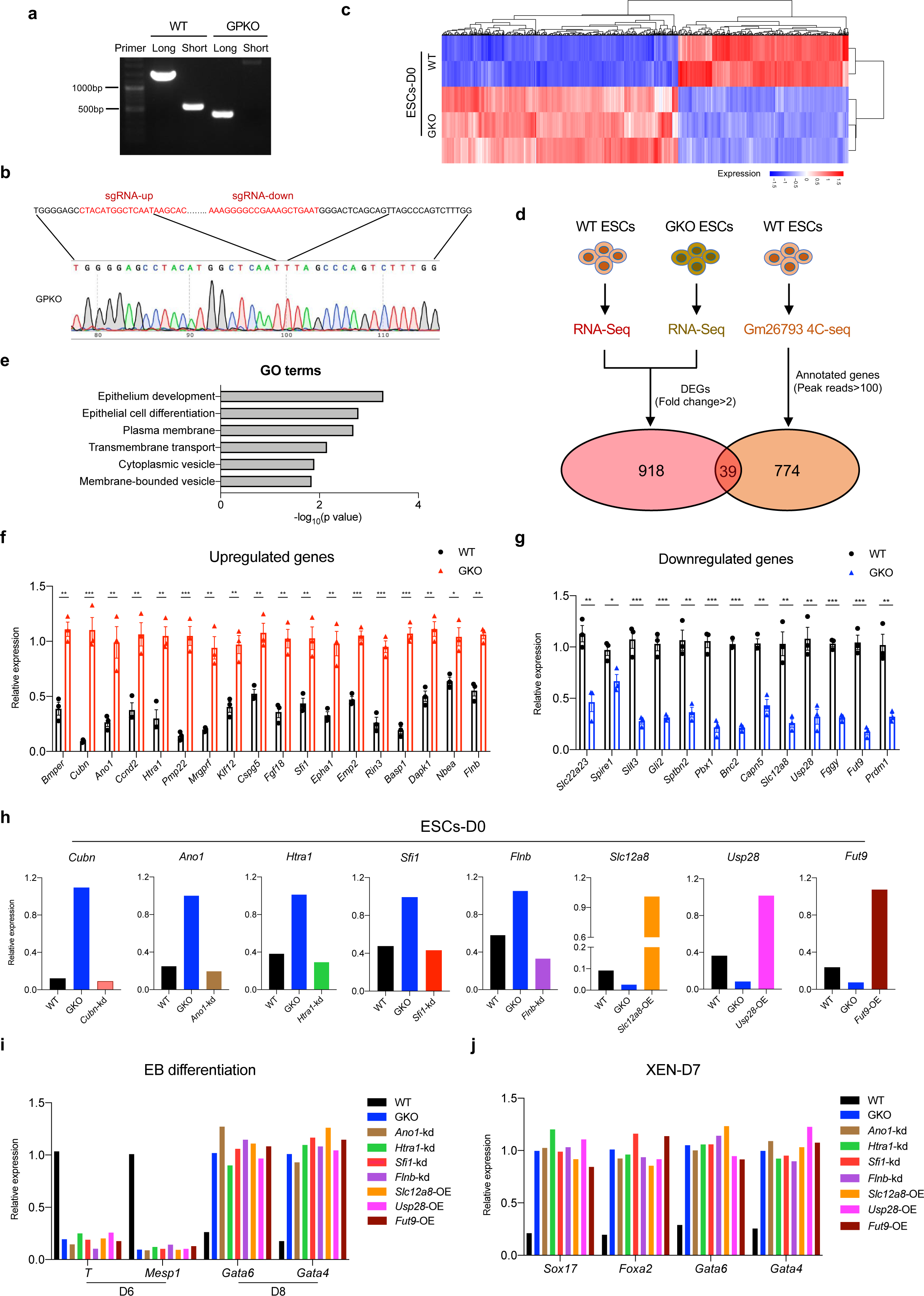
Functional screening of genes regulated by *Gm26793* locus. (a) DNA electrophoresis result demonstrating the removal of promoter region of *Gm26793* in mESCs. (b) The genomic sequence of GPKO was confirmed by Sanger sequencing. (c) Heatmap highlighting the direct transcriptomic comparison between WT and GKO mESCs. (d) Venn diagram showing the workflow for the screening of potential *Gm26793* targets by the integrative analyses of RNA-seq and 4C-seq data. (e) GO term enrichment of the selected 39 interaction genes. (f-g) qPCR validation of the top 31 genes in 39 selected candidates, in which 18 genes were up-regulated (d) and 13 genes were down-regulated (e) in GKO mESCs. Data were shown as means ± SEM. Student’s *t*-test, \**p*<0.05, ** *p*<0.01, *** *p*<0.001. (h) Perturbation of candidate genes through knockdown or overexpression in GKO mESCs. (i-j) qPCR analyses of lineage-specific genes in EB (g) and XEN (h) differentiation after expression perturbation of the candidate genes.

**Extended Data Fig 6.**
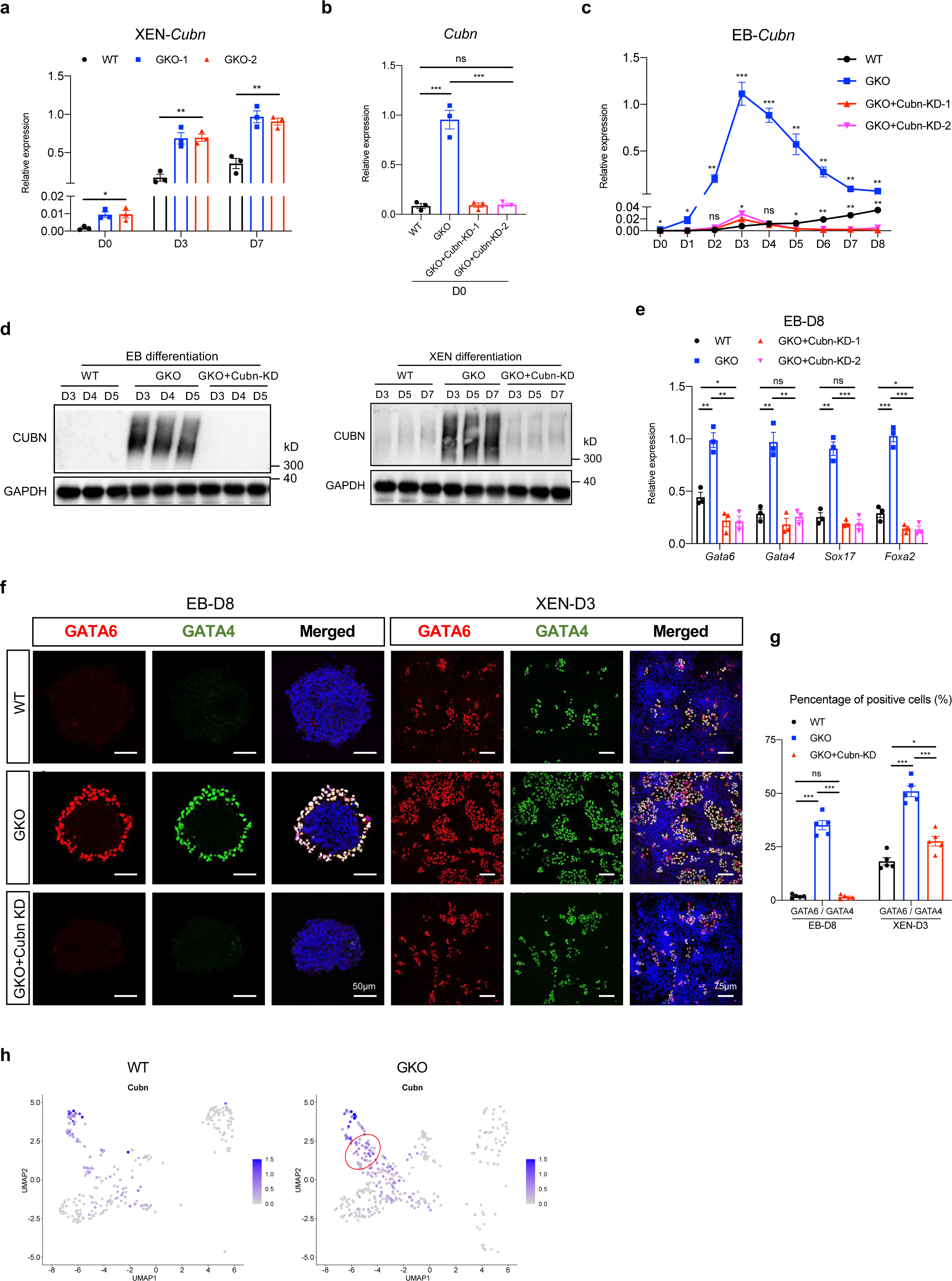
*Cubn* knockdown recovers the enhanced differentiation potential of PrE in GKO cells. (a) Constant activation of *Cubn* in GKO cells during XEN differentiation. (b) Bar plot showing the successful knockdown of *Cubn* expression in GKO+Cubn KD mESCs. (c) Relative expression dynamics of *Cubn* in indicated cell groups during EB differentiation. (d) Protein expression changes of CUBN during EB (left) and XEN (right) differentiation upon *Cubn* knockdown. (e) Reduced expression of PrE-related genes on day 8 of EB differentiation upon *Cubn* knockdown. (f-g) Immunostaining images reporting the rescue of GATA6 and GATA4 expression pattern after *Cubn* knockdown in GKO cells. The specific differentiation system and timepoint for sampling were indicated at the top of each image. Scale bar, 50 μm and 75 μm. Quantification of the data was shown in (g). (h) UMAP plot showing the up-regulation of *Cubn* in transient epiblast and primitive endoderm cells. All data were shown as means ± SEM. Student’s *t* test, \**p*<0.05, ** *p*<0.01, *** *p*<0.001.

**Extended Data Fig 7.**
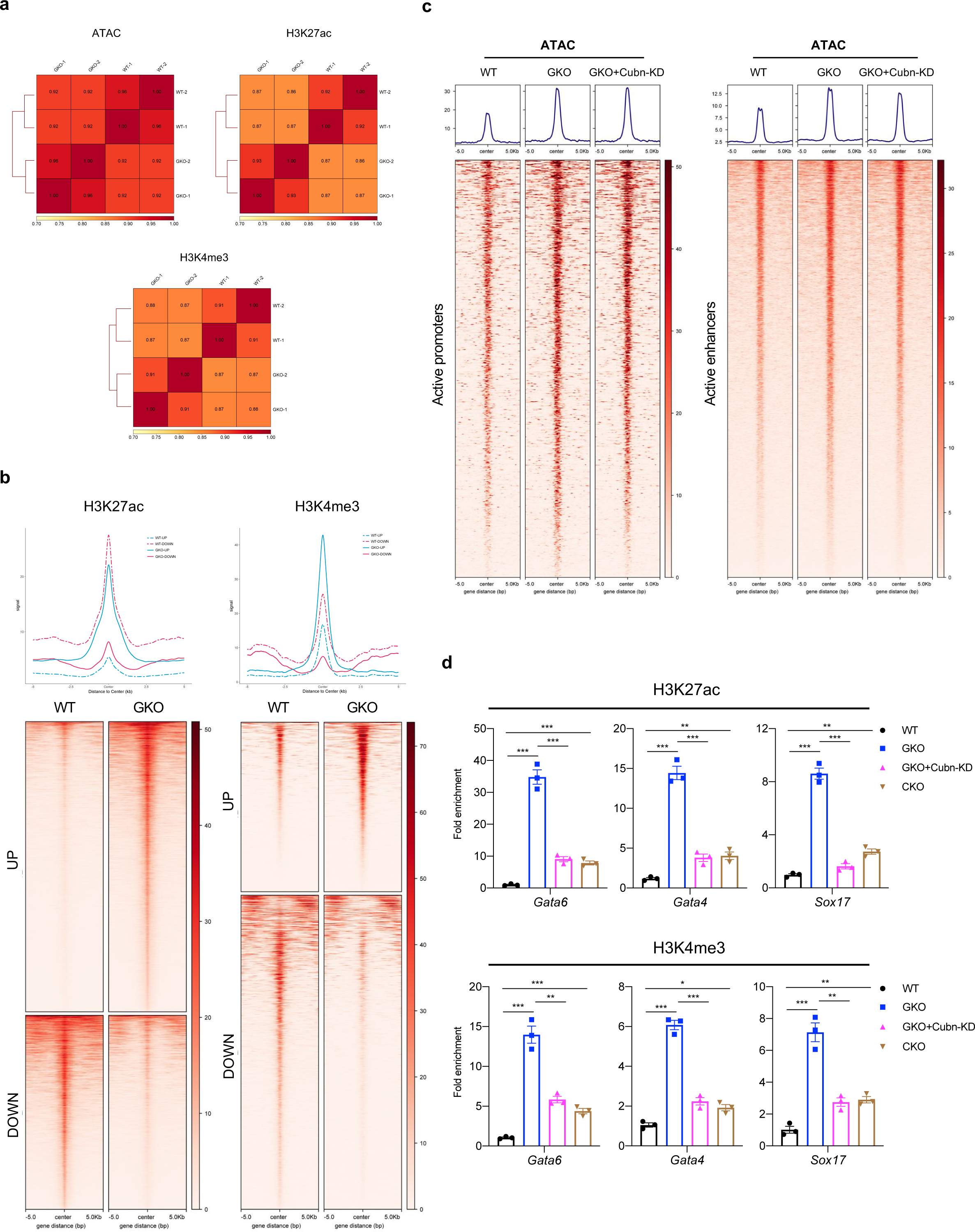
ATAC-seq and ChIP-seq analysis. (a) Spearman correlation of ATAC-seq, H3K27ac, as well as H3K4me3 signal in both WT and GKO replicates. (b) Heat maps showing the global alteration of H3K27ac and H3K4me3 distribution in GKO cells in compare with WT mESCs. The averaged signal intensities for respective histone modification in each cell group were plotted at the top of each heatmap. (c) Global distribution of ATAC-seq signal around selected genomic regions in WT, GKO, and GKO+Cubn-KD cells. (d) ChIP-qPCR analyses validating the dynamic changes of H3K27ac and H3K4me3 enrichment around selected gene loci in indicated mESCs. Data were shown as means ± SEM. Student’s *t* test, \**p*<0.05, ** *p*<0.01, *** *p*<0.001.

**Extended Data Fig 8.**
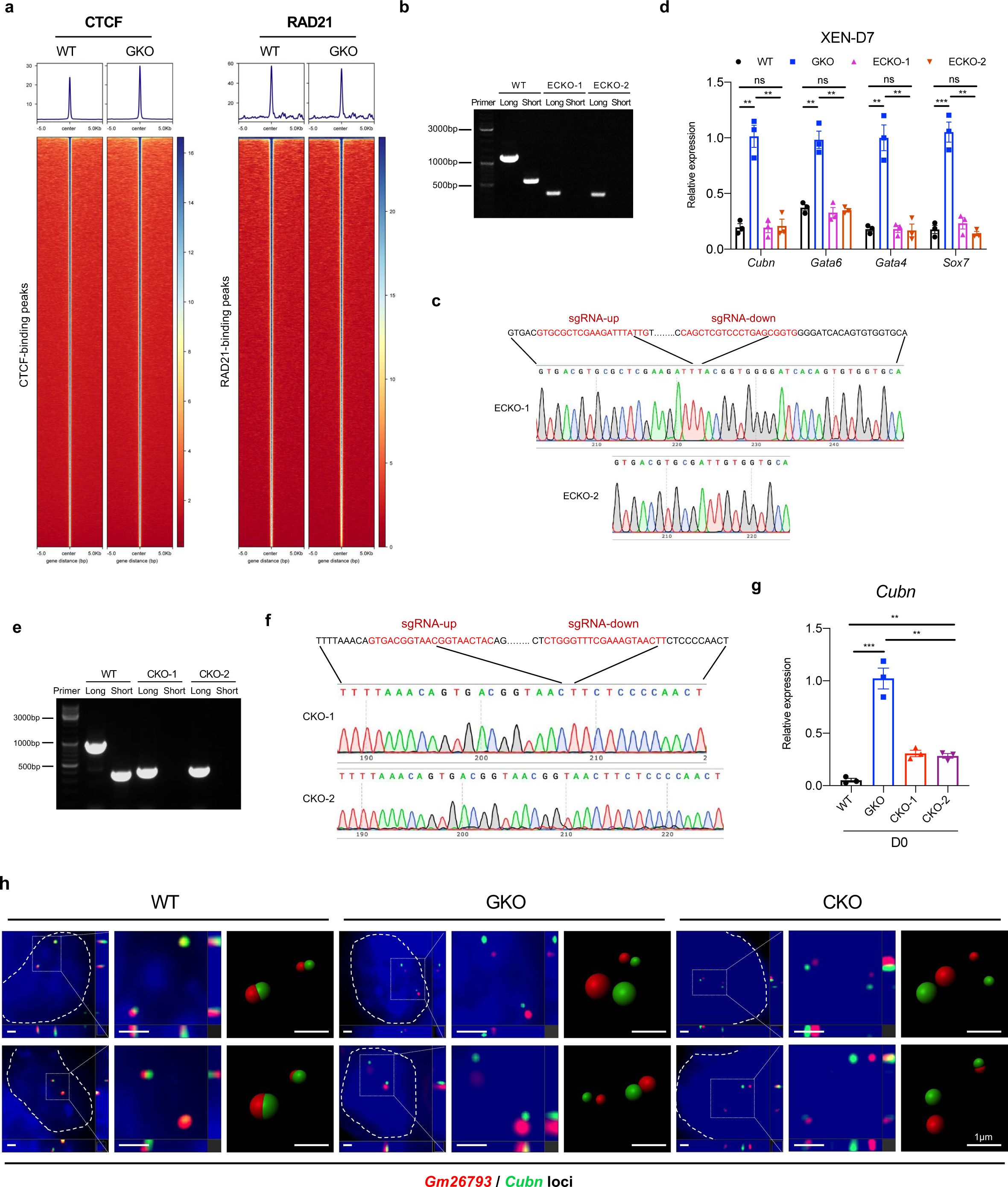
Establishment and functional validation of ECKO and CKO mESCs. (a) Heatmaps showing global CTCF and RAD21 binding signals in WT and GKO mESCs. (b-c) DNA electrophoresis showing the efficient removal of ECKO locus in mESCs. The acquired truncated DNA sequences were determined by sanger sequencing (c). (d) Bar plot showing the relative expression level of PrE-related marker genes in WT, GKO, and ECKO XEN. (e-f) DNA electrophoresis showing the efficient removal of CKO locus in mESCs. The acquired truncated DNA sequences were determined by sanger sequencing (f). (g) qPCR analyses identifying the upregulation of *Cubn* expression in CKO mESCs. (h) Representative images showing 3D-reconstruction of *Gm26793* (red) and *Cubn* (green) loci (two alleles) in indicated mESCs. Scale bar, 1 μm. All data were shown as means ± SEM. Student’s *t* test, \**p*<0.05, ** *p*<0.01, *** *p*<0.001.

## Methods

### Mouse strains

All mice were housed in individually ventilated cages (IVC) under specific pathogen-free conditions and handled according to the guidelines of the Animal Ethical Committee of the Institute of Biochemistry and Cell Biology, Shanghai Institutes for Biological Sciences, Chinese Academy of Sciences. To generate GKO mice, we firstly derived GKO DKO-AG-haESCs by CRISPR-Cas9. Then GKO female mice were constructed via intracytoplasmic AG-haESCs injection (ICAHCI) ^105^ and bred with wild type (WT) males (C57BL/6J) to produce GKO heterozygous offspring. The heterozygous mice were mated internally to generate GKO homozygous mice. WT embryos were collected from the C57BL/6J background mice.

### Embryos

For preimplantation embryos, all zygotes were obtained from superovulated and fertilized female mice, and then cultured in EmbryoMax Advanced potassium-supplemented simplex optimized medium (KSOM) with amino acids under mineral oil on polystyrene plates. Embryos were maintained in a humidified incubator at 37°C with 5% CO2 until blastocyst stage. For gastrula, embryos were removed from the implantation site as described previously ^106^. Brieflly, plugged female mice were picked after mating and counted as embryonic day 0.5 (E0.5). Mice were euthanized when embryos developed into nominal day 7.5. The embryos were acquired through removal of the surrounding decidua and Reichert’s membrane by using sharpened surgeon tweezers.

### mESCs culture and differentiation

mESCs (E14) were cultured under feeder-free conditions on gelatinized dishes in DMEM medium supplemented with 15% FBS, 1% GlutaMAX, 1% NEAA, 1 mM sodium pyruvate, 0.1 mM β-mercaptoethanol, 1% penicillin/streptomycin, 1000 U/ml mouse LIF, 3 μM CHIR99021, 1 μM PD0325901, and passaged by single-cell trypsinization every 2-3 days. For EBs differentiation, mESCs were dissociated with 0.05% trypsin and suspended in differentiation medium consisting of DMEM, 10% FBS, 1% GlutaMAX, 1% NEAA, 1 mM sodium pyruvate, 0.1 mM β-mercaptoethanol and 1% penicillin/streptomycin. 1X10^5^ cells/ml were plated in Petri-dishes and cultured for 8 days. Every 2 days, we need to change the medium and divide the differentiated EBs into fresh Petri-dishes. For XEN differentiation, 1X10^4^ cells/cm^2^ were seeded on gelatin-coated dishes and cultured in standard XEN medium consisting of RPMI-1640, 15% FBS, 1% GlutaMAX, 1% penicillin/streptomycin and 0.1 mM β-mercaptoethanol for 1 day, then the medium was changed to derivation XEN medium (standard XEN medium supplemented with 0.01 μM RA and 10 ng/ml Activin A). After two days of culture, differentiated cells were dissociated into single cells and plated at 1:1 ratio on MEF-coated dishes, hereafter maintained in standard XEN medium.

### CRISPR-Cas9-mediated knockout

The deletion of *Gm26793* locus and CTCF binding sites in mESCs were performed by CRISPR-Cas9 gene editing system. Briefly, a pair of sgRNAs (upstream and downstream) were designed and inserted into pX330-mCherry vector. mESCs were transfected with 5 μg sg-up&down plasmids by Lipofectamine^TM^ 2000 and cultured for 24h. 1X10^4^ mCherry-positive single cells were sorted to seed on a 10cm gelatinized dish. After 4-6 days of culture, individual colonies were picked up and expanded in 48-well plates. Finally, genomic deletion mESCs were validated by PCR and Sanger-sequencing. sgRNA oligos and genotyping primers were presented in Supplementary Table 11 and 12.

### RNAi and overexpression assays

For knockdown experiments, shRNAs were constructed into lentiviral vector pLKO.1 for lentiviral packaging. After 48 hr transfection into mESCs, puromycin resistant cells were selected for testing the efficiency of knockdown. The sequence of shRNAs targeting specific genes were designed by using online tool GPP Web Portal (https://portals.broadinstitute.org/gpp/public/) and listed in Supplementary Table 12. For overexpression experiments, ORFs were similarly cloned into lentiviral vector Fugw-IRES-dsRed or Fuw-TRE-P2A-mCherry (inducible). RFP-positive mESCs were eventually sorted for functional analysis. The inducible overexpression system will proceed under the treatment of Dox.

### Whole-mount in situ hybridization

Digoxigenin (DIG)-labeled riboprobes were synthesized as previously reported ^107^. Primers used for amplifying probe templates were listed in Supplementary Table 11. In brief, E7.5 embryos were fixed with 4% PFA, dehydrated and rehydrated through 100%, 75%, 50% and 25% methanol. Samples were then treated with 10 μg/ml proteinase K for 10 min and post-fixed with 0.1% glutaraldehyde for 30 min. Approximately 1 μg/ml DIG-labeled RNA-probe was incubated with the embryos at 70℃ overnight. After washing, the embryos were incubated in anti-DIG-AP at 4℃ overnight, then washed and stained with NBT and BCIP for imaging.

### Quantitative real-time PCR analysis (qPCR)

Total RNA was extracted from cultured cells by using TRIZOL reagent and 500 ng to 2 μg RNA was reversely transcribed into first-strand cDNA by FastKing RT Kit (Tiangen, KR116). qPCR analysis was performed with Mastercycler Realplex2 (Eppendorf) using Stormstar SYBR green qPCR master mix (DBI-2144). The relative expression of target genes was normalized to internal control *Gapdh* and quantified by 2^-ΔΔCt^ methods. qPCR primers for specific genes were presented in Supplementary Table 11.

### Immunostaining

After fixation with 4% paraformaldehyde (PFA) for 30 min, the cultured cells were permeabilized and blocked with 0.3% Triton X-100/5% BSA in PBS for 1 hr at room temperature. Then, the samples were incubated with primary antibodies (1:200) at 4℃ overnight. The next day, samples were washed 3 times and incubated with fluorescence-conjugated secondary antibodies (1:500-1:1000) for 1 hr at room temperature. The nuclei were stained with DAPI (1:1000). For staining of embryoid bodies, samples were dehydrated in 20% sucrose at 4℃ overnight after fixation, and then embedded in OCT for cryosection. Slides immunostaining were conducted as described above. The antibody information was listed in Supplementary Table 13 and images were taken from Leica TCS SP8 confocal laser-scanning microscope.

### Western blotting

The harvested cells were lysed in RIPA buffer with protease and phosphatase inhibitors for 30 min on ice. After centrifugation, proteins in supernatant were quantified and added to loading buffer for heating 10 min at 100℃. 20 ug total protein were separated by SDS-PAGE and transferred to PVDF membranes. The membranes were blocked with 5% BSA and incubated with primary antibodies (1:1000) at 4℃ overnight. After three times wash with TBST, the membranes were incubated with HRP-conjugated secondary antibodies (1:2000) for 1 hr at room temperature, and the target proteins were detected by SuperSignal™ West Pico PLUS Substrate (Thermo Scientific, 34580) subsequently.

### Geo-seq of embryoid body samples

Geographical position sequencing (Geo-seq) was performed as previously described ^69^. Briefly, whole embryoid bobies were embedded in OCT and cryosectioned at a thickness of 20 μm. Sections were mounted on polyethylene-terephthalate-coated slides, fixed with ethanol and stained with 1% DAPI in 75% ethanol solution. Then, approximately 20 cells in designated region were captured by laser microdissection (MMI Cellcut Plus system) and lysed in 50 μl 4M guanidine isothiocyanate for 15 min at 42℃. After isolation through ethanol precipitation, dissolved RNA was immediately reversely transcribed into cDNA and amplified by Smart-seq2.

### CRISPR-dCas9-based imaging

To visualize *Gm26793*-*Cubn* loci interaction, we used the CRISPR-mediated DNA labeling system to achieve non-repetitive DNA imaging, according to previous description ^93^. Briefly, 30bp crRNAs targeting *Gm26793* and *Cubn* loci were synthesized with fluorescent labeling (Cy5-Gm26793; TAMRA-Cubn) at the 5′-end (Sango). The sequences of crRNAs used in this study were presented in Supplementary Table 12. The crRNAs and tracrRNA were annealed and incubated with dCas9 protein (IDT) to form fluorescent RNA protein complexes (fRNPs). Then, mESCs were transfected with the pre-assembled fRNP pool by electroporation (program: OP315 or CD112) using an SE Cell line 4D-Nucleofector™ X kit (Lonza, Catalog#: V4XC-1024). The electroporated cells were plated in Nunc Glass Bottom Dishes (Thermo Fisher, 150680) and cultured for 12-24 h before imaging. Microscopic imaging was performed on a Leica DMi8 Inverted Microscope using sCMOS camera and APO ×63/1.4 oil objective or a Leica TCS SP8 STED equipped with the spectral flexibility of WLL for excitation and an HC PL APO ×100/1.4 oil objective with Z stacks from 0.27 to 5 μm. Nuclei were visualized using DAPI for fixed cells or NucBlue™ Live ReadyProbes™ (Thermo Fisher, R37605) for living cells.

### Image analysis

To quantify the 3D distance between *Gm26793* and *Cubn* loci, the imaging data were processed using deconvolution wizard and chromatic Aberration Corrector to generate the ICS2 files after adjusting imaging parameters of channel (excitation and emission wavelengths), type of microscope, material of vehicle and imaging optical path media, automatically generated theoretical PSF and correct Z-drift in Huygens software. The ICS2 files were loaded by Imaris to measure the distance between *Gm26793*-*Cubn* loci after performing spot simulation based on each fluorescence channel maximum value corresponding to the *Gm26793*-*Cubn* loci.

### Bulk RNA-seq data processing and analysis

Data quality control was performed to ensure the reliability of the results. Quality control metrics such as sequence quality scores, GC content, and adapter content were assessed using FastQC (v0.11.9). Low-quality reads and adapters were trimmed or removed. Then, the high-quality reads were aligned to a reference genome Hisat2 (v2.2.1) and the reference genome used was mm10. Gene-level expression quantification was performed using featureCounts (v1.5.3). This step assigns reads to genes and generates a count matrix representing the number of reads mapped to each gene in each sample. The count matrix was normalized to account for differences in sequencing depth and gene length as fragments per kilobase of transcript per million mapped reads (FPKM). Differential expression analysis was performed to identify genes that were differentially expressed between conditions or groups of interest using the DESeq2 package with count matrix as input.

### Annotation and identification of differentially expressed lncRNAs during mouse gastrulation

Raw lncRNA-seq data was obtained from GEO-seq datasets ^63^. The high-quality reads were aligned to a reference genome using a read aligner Tophat2. The reference genome used was mm10. An annotation file containing known lncRNA transcripts was obtained from databases such as GENCODE. The alignment results were filtered to retain only reads that mapped to the annotated lncRNA regions. Quantification of lncRNA expression levels was performed with Cufflinks. This step assigns reads to lncRNA transcripts and generates a FPKM matrix. Differentially expressed lncRNAs (DELs) were identified as follows: (1) calculation of the variance of each expressed lncRNA across all samples and selection of top approximately 1000 genes as highly variable genes; (2) hierarchical clustering with correlation distance metric based on z-score normalized expression of highly variable genes to identify preliminary domains according to distinctly separated dendrogram; (3) identification of the inter-domain DELs, on the basis of expression of highly variable genes by pairwise comparisons of preliminary domains using t-test (P < 0.05) and fold change (FC >= 1.5); (4) combination of top highest and lowest principal component (PC)-loading genes (by using FactoMineR (2.8) in R) from several selected significant PCs by jackstraw to identify the DELs (top 300 genes for each of PC1– 4). Finally, Kmeans clustering was applied to determine the final spatial domains of embryo based on the expression profile of DELs, and BIC-SKmeans algorithm was applied to determine the optimal number of gene groups and perform gene clustering analysis based on the z-score normalized expression profile of DELs. The clustering heatmap was visualized through ComplexHeatmap (2.15.1).

### Weighted gene co-expression network analysis (WGCNA)

Co-expression networks were constructed using WGCNA (v 1.72.1) package in R ^68^. Firstly, we created a matrix of pairwise correlations between all pairs of genes across the measured samples. Next, we identified the soft thresholding power (β) value based on the scale-free topology network criterion and converted the expression matrix into a adjacency matrix. The adjacency matrix was then transformed into a topological overlap matrix (TOM) to capture the interconnectedness of genes within the network. Next, the topological overlap dissimilarity was calculated using TOM, followed by hierarchical clustering to identify separated gene modules. A dynamic tree cutting algorithm was employed for gene module determination with a minimum size of 30 and highly similar modules would be merged automatically. Then, the module eigengene (ME), which represents the first principal component of each module, was estimated and summarizes the overall expression pattern of genes within one module. Finally, we can perform module-trait relationship analysis to assess the correlation coefficient between module eigengenes and sample traits or phenotypes of interest. This analysis helps to identify certain unique gene modules associated with specific biological conditions.

### 4C-seq analysis

Circle chromosome conformation capture (4C) was carried out by a modified published protocol ^108^. In brief, 5X10^6^ mESCs were fixed in 1% formaldehyde solution and quenched with 0.125 M glycine, then rinsed by DPBS twice before frozen in liquid nitrogen. After thawing on ice, fixed cells were resuspended in lysis buffer (50mM Tris-HCl pH 7.5, 0.5% NP-40, 1% Triton X-100, 150mM NaCl, 5mM EDTA) and 0.5% SDS buffer for 30 min, respectively. Each sample was then permeabilized in 1% Triton X-100 and digested with DpnII for 3 hr at 37°C. After first ligation, fragmented DNA was purified by using phenol-chloroform and ethanol precipitation upon treatment with RNase A and Proteinase K, then digested again with Csp6I for 3 hr at 37°C. After second ligation, circularized DNA was purified with AMPure XP beads and 4C-libraries were finally generated by two rounds of PCR and purified by QIAGEN column. First PCR step was conducted to reversely amplify the DNA fragments ligated to the *Gm26793* viewpoint. Second PCR step was to add Illumina index for high-throughput sequencing. Primers used for 4C library construction were presented in Supplementary Table 12. Proteinase inhibitor cocktail and PMSF required to be added before the second digestion to prevent protein degradation. Three biological replicates were performed for 4C-seq analysis of *Gm26793* locus. Sequencing reads with 5’end matching the inverse PCR primer sequence were selected and trimmed, remaining sequences containing DpnII and Csp6I sites were mapped to mm10 assembly using Bowtie2 (v2.4.5) and the interaction regions were identified on the basis of pipeline proposed by Krijger et al.

### ChIP-seq analysis

Chromatin immunoprecipitation (ChIP) was performed as previously described ^109^. Briefly, cross-linked mESCs were lysed in lysis buffer and fragmented to a size range of 200-500 bp by using Bioruptor Pico. Then, solubilized fragmented chromatin was immunoprecipitated with primary antibodies (CTCF, RAD21, H3K27ac and H3K4m3) and pulled down by protein G beads. Reverse crosslink was performed at 65℃ for at least 4 hours. Subsequently, ChIP-DNA was treated with RNase A and Proteinase K, precipitated with ethanol and dissolved in nuclease free water. Proteinase inhibitor cocktail and PMSF were added in all IP assays to inhibit protein degradation. Additionally, 20mM sodium butyrate was added in H3K27ac group to inhibit histone deacetylase. Finally, ChIP libraries were prepared by using NEBNext® Ultra™ DNA Library Prep Kit (NEB, E7770L) for Illumina. ChIP-seq reads were mapped to mm10 genome with Bowtie2 (v2.4.5) using default parameters, then the peaks were called using MACS2 (v2.2.7) to identify regions of the genome that exhibit significant enrichment for CTCF, RAD21 and histone modifications compared to background. Differential peaks of ChIP-seq experiments were called with the R package, DiffBind (v3.6.5), under default settings. Heatmaps of ChIP-seq signal enrichment were generated by the Python package, deepTools (v3.5.1). Annotation of ChIP-seq peaks was done by ChIPseeker (v1.32.1).

### ATAC-seq analysis

For ATAC-seq, 5X10^4^ cells were harvested and resuspended in lysis buffer (10mM Tris-HCl pH 7.4, 0.15% NP-40, 10mM NaCl, 3mM MgCl2). After vortex for 3 times every 3 min, the tube was centrifuged at 1000 g for 10 min at 4°C and the supernatant was discarded. Then, the cell pellet was resuspended in fragmentation buffer (5XTTBL, TTE V50 Mix) (Vazyme, TD501) and incubated at 37°C for 30 min. Immediately, fragmented DNA was purified by QIAGEN MinElute Kit and sequencing libraries were generated by PCR using NEB Q5 Master mix (E7649A). The size of library was selected and purified with AMPure XP beads. The package versions involved in ATAC-seq data processing are the same as ChIP-seq analysis.

### Blastocyst collection and single-cell isolation

To obtain adequate blastocysts, WT and GKO female mice (8-10 weeks) were induced to superovulate by injection of 7.5 IU pregnant mare’s serum gonadotropin (PMSG) followed by 7.5 IU human chorionic gonadotropin (hCG), and then were mated with male mice. Vaginal plugs were checked the next morning. Zygotes were collected from the oviduct 13-15 hrs after hCG injection. Embryos were cultured with KSOM medium containing amino acids in an incubator with 5% CO2 at 37°C. Early and late blastocysts were collected at E3.5 and E4.5, respectively. Single-cell isolation of blastocysts was performed as follow. After removal of zona pellucida using acid Tyrode solution, the embryos were dissociated with 1% trypsin/EDTA for 30-40min at 37°C, then washed twice and resuspended in 0.1% BSA/PBS. Single cells were manually picked into PCR tubes by mouth pipette under a microscope.

### scRNA-seq analysis

Single cells were lysed in lysis buffer containing 0.45% NP-40, followed by reverse transcription using SuperScript II reverse transcriptase. Subsequently, the entire cDNA was amplified with 2×KAPA Mix and PCR products were purified with AMPure XP beads and quantified with Qubit. cDNA libraries were then constructed with TruePrep DNA Library Prep Kit V2 (Vazyme, TD503) for Illumina. In total, 1007 single cells were sequenced for further analysis. Library construction and purification were completed on Agilent Bravo automatic liquid-handling platform. Raw counts of genes were calculated for each cell using the same workflow as for bulk RNA-seq. Seurat (v4.3.0) was utilized for the analysis of single-cell data. Mapping and annotation of query datasets were performed referring to this workflow (https://satijalab.org/seurat/articles/integration_mapping.html). Briefly, UMAP were applied to visualize the cell clusters in two dimensions. Clustering was performed based on the shared nearest neighbor (SNN) graph using algorithms such as Louvain clustering or density-based clustering. Cell type annotation was performed by comparing the cluster-specific marker genes with known cell type marker genes from databases or literature. Functional enrichment analysis, such as Gene Ontology (GO) or pathway enrichment, was performed with Metascape to identify the biological processes, molecular functions and pathways enriched in specific cell clusters. Monocle 3 ^81^ (1.2.9) was applied to reconstruct the pseudotime trajectories. We manually selected Inner cell mass as root node and used variable genes identified in Seurat as ordering genes in the Monocle 3 pipeline. Monocle 3 used a graph embedded algorithm to learn a trajectory that fit the UMAP coordinated cell clusters.

### Quantification and statistical analysis

For quantification of the immunostaining, we counted the distribution of corresponding antibody-positive cells in five randomly selected visual fields. All experiments were performed with at least three biological replicates. All error bars were presented as mean ± SEM. The results were analysed using two-tailed unpaired *t*-test, and *P* < 0.05 was considered statistically significant at the 95% confidence level. GraphPad Prism 8 and Excel were used for statistical calculations and generation of plots.

### Data and code availability

The raw sequence data reported in this paper have been deposited in the Genome Sequence Archive in National Genomics Data Center, China National Center for Bioinformation/Beijing Institute of Genomics, Chinese Academy of Sciences (GSA: CRA011812, CRA011818, CRA011832, CRA011839, CRA011986) that are publicly accessible at https://ngdc.cncb.ac.cn/gsa.

## Acknowledgements

This work was supported in part by the National Key Basic Research and Development Program of China (2019YFA0801402, 2018YFA0800100), the Strategic Priority Research Program of the Chinese Academy of Sciences (XDA16020308), National Natural Science Foundation of China (32130030, 31900454).

## Author contributions

Z.L., X.Y. and N.J. conceived the study. X.Y., D.Z., J.L. and N.J. supervised the project. Z.L. and X.Y. designed and performed the experiments. J.C. and Y.F. conducted the bioinformatic analyses. Y.M. and Y.C. constructed the *Gm26793* knockout mice and collected single cells from mouse blastocysts. X.W. performed the dCas9-mediated DNA imaging. Y.C., Y.Q., and M.W. helped to breed *Gm26793* knockout mice and complete scRNA-seq. Z.L., X.Y. and N.J. wrote the paper with the help from all other authors.

## Declaration of interests

The authors declare no competing interests.

## Notes

### Competing Interest Statement

The authors have declared no competing interest.

